# Tumor cell-intrinsic stress states drive sensitivity to CAR T cell-therapy in pancreatic cancer

**DOI:** 10.1101/2025.07.24.666408

**Authors:** Julia Fröse, Evelyn Chen, Charles A. Whittaker, Paul Leclerc, Sean Doherty, Adam Langenbucher, Jasmine Shao, Riley D. Hellinger, Daniel Goulet, Tuomas Tammela, Francisco J. Sánchez-Rivera, Michael T. Hemann

**Affiliations:** The David H. Koch Institute for Integrative Cancer Research; Cambridge, MA 02142, United States; Department of Biology, Massachusetts Institute of Technology; Cambridge, MA 02142, United States; Cancer Biology and Genetics Program, Sloan Kettering Institute, Memorial Sloan Kettering Cancer Center, New York, New York; David M. Rubenstein Center for Pancreatic Cancer, Memorial Sloan Kettering Cancer Center, New York, New York

## Abstract

Chimeric antigen receptor (CAR) T cell therapy has transformed the treatment of hematologic cancers, but has shown limited efficacy in solid tumors, including pancreatic ductal adenocarcinoma (PDAC). The cellular and molecular factors that influence CAR T cell therapy response remain largely unknown. By integrating modular *in vivo* CRISPR screens with single cell genomics and immunocompetent orthotopic models of PDAC, we uncover oxidative and proteotoxic stress pathways as previously unknown tumor-intrinsic modulators of CAR T cell therapy response. Disruption of stress-regulatory genes, particularly *Keap1* and *Slc33a1*, sensitizes PDAC tumors to CAR T cell killing *in vivo*. Mechanistically, hyperactivation of the *Nrf2* pathway by genetic ablation of the *Keap1* tumor suppressor or endogenous engineering of a clinically observed *Keap1* mutation enhances tumor susceptibility to CAR T cell therapy. Thus, tumor-intrinsic molecular stress phenotypes accompanying malignant tumor progression can induce unexpected cell state-specific vulnerabilities to cell-based immunotherapies. These findings provide a mechanistic molecular foundation to improve the efficacy of CAR T cell therapy in solid malignancies and to better stratify cancer patients by tumor genotype.

**Statement of significance:** CAR T cell therapy remains an unsolved challenge for pancreatic cancer. Factors that influence CAR-T cell therapy response remain largely unknown, in part due to lack of robust and scalable experimental models. By integrating in vivo genetic screens with single cell genomics in an orthotopic, immunocompetent model of pancreatic cancer, we uncover cell-intrinsic stress states as key regulators of CAR T cell response.

## Introduction

Pancreatic ductal adenocarcinoma (PDAC) is projected to become the second leading cause of cancer death by 2030^1,2^. PDAC lacks effective treatment options, and innovative strategies for therapeutic intervention are urgently needed^3^. The use of chimeric antigen receptor (CAR) T cell therapy, in which a patient’s own T cells are engineered to recognize and kill cancer cells, has shown remarkable success in hematological malignancies in recent years^4–6^. The adaptation of these therapies to solid tumors such as pancreatic cancer holds great promise^7^. However, clinical trials for CAR T cell therapy in PDAC have shown limited efficacy in patients^8^. To date, no universal mechanism underlying the therapeutic success or failure has been identified.

CAR T cells face a host of challenges, especially in solid tumor microenvironments, such as inefficient homing to the tumor site, immunosuppressive tumor microenvironments, and lack of persistence in the tumor due to exhaustion or other factors^9–11^. Many strategies for improving CAR T cell therapy efficacy have focused on further engineering CAR T cells to improve their persistence and expansion or prevent exhaustion in cancer patients^12–17^. However, initial studies examining tumor response to immune checkpoint blockade have shown that tumor cell-intrinsic properties can also play important roles in influencing the response to immunotherapy^18^. Recent studies have identified tumor-intrinsic mechanisms of CAR T cell therapy resistance and disease relapse, including antigen loss^19–21^, impaired death receptor signaling^22^, and alterations in the IFN-γ pathway^23–26^. Nevertheless, to date, no cancer-specific dependencies have been discovered that can influence CAR T cell therapy response. It also remains unclear whether the cancer alterations that underlie tumor responses to conventional or targeted chemotherapy can similarly impact response to CAR-T treatment.

Solid tumors are complex ecosystems of tumor cells, as well as multiple types of infiltrating immune cells, fibroblasts, and endothelial cells^27^. PDAC tumors are particularly complex malignancies characterized by substantial inter- and intertumoral heterogeneity, as well as an unusually dense fibrotic stroma^28,29^. Recent studies have also shown that PDAC tumors harbor distinct phenotypic states, including classical, basal, and mesenchymal states^30–32^. Additionally, different cell states have been shown to modulate response to targeted therapies and cytotoxic T cell killing^33,34^. It is unclear whether there are any tumor cell states in PDAC that are particularly sensitive to CAR T cell treatment.

A number of studies focused on CAR T cell therapy resistance in solid tumors have relied on CRISPR-Cas9 knockout (KO) screens in cell culture, immunodeficient mouse models, or heterotopic locations ^26,35^. While these screens have uncovered important biology, *in vivo* screening in orthotopic locations and in the presence of a fully functional immune system may reveal previously unknown tumor-intrinsic mediators of CAR T cell response. Understanding which tumor-intrinsic mechanisms mediate resistance to immunotherapy in relevant physiological settings is key in designing effective treatments for patients.

To uncover tumor cell-intrinsic mechanisms of resistance to CAR T cell therapy in PDAC, we performed iterative CRISPR-Cas9-based pooled KO screens in a fully immunocompetent, orthotopic model of PDAC treated with CAR T cells. In this physiological setting, we uncovered previously unknown modulators of CAR T cell therapy response. Importantly, single-cell RNA sequencing (scRNA seq) of CAR T cell-treated PDAC tumors revealed that therapy-resistant tumor cells exhibit a metabolic cell state characterized by reduced antioxidant pathway activation, reactive oxygen species (ROS) signaling, and glutathione metabolism driven by the oxidative stress response master regulator Nrf2. Using these two orthogonal approaches, we show that loss of genes involved in proteotoxic and oxidative stress, particularly *Slc33a1* and *Keap1*, is sufficient to sensitize PDAC tumors to CAR T cell treatment *in vivo*. Intriguingly, hyperactivation of the Keap1/Nrf2 antioxidant response pathway by genetic ablation of *Keap1* or endogenous engineering of a tumor-derived *Keap1* mutation is sufficient to sensitize PDAC tumors to CAR T cell therapy. This work presents a general strategy to integrate functional genomics with pre-clinical mouse models to study mechanisms of response and resistance to state-of-the-art cell-based immunotherapies. More broadly, our data indicate a potential avenue for therapeutic synergy by targeting pathways involved in cellular stress states to increase the efficacy of CAR T cell therapy in PDAC.

## Results

### A scalable mouse model of PDAC to study CAR T cell therapy resistance

To identify genetic mediators CAR T cell therapy response in an unbiased manner, we established a murine PDAC model amenable to large-scale CRISPR KO screens and targeting by murine CAR T cells. For this purpose, a PDAC cell line derived from autochthonous pancreatic tumors in the KPC (*LSL-Kras^G12D/+^;LSL-Trp53^R172H/+^;Pdx1-Cre*) mouse model (**Figure 1A**) was engineered to express both *firefly luciferase* (Luc) and *SpCas9*^36^. From this population, a Cas9-expressing (Cas9+) PDAC clone with high genome editing efficiency (**Figure S1A-B**) was isolated. This clone was further modified to express human CD19 (hCD19) without the intracellular domain, as an antigen for CAR T cell treatment (hCD19+ PDAC; **Figure S1C**). We selected hCD19 as a tumor antigen for three reasons. First, it allowed us to use established anti-hCD19 CAR constructs. Second, it narrowed the potential scope of biology emerging from the screen results by eliminating the relevance of genes involved in endogenous antigen presentation. Third, this strategy can be extrapolated to other tumor types that may lack suitable murine CAR T cell targets.

**Figure 1:**
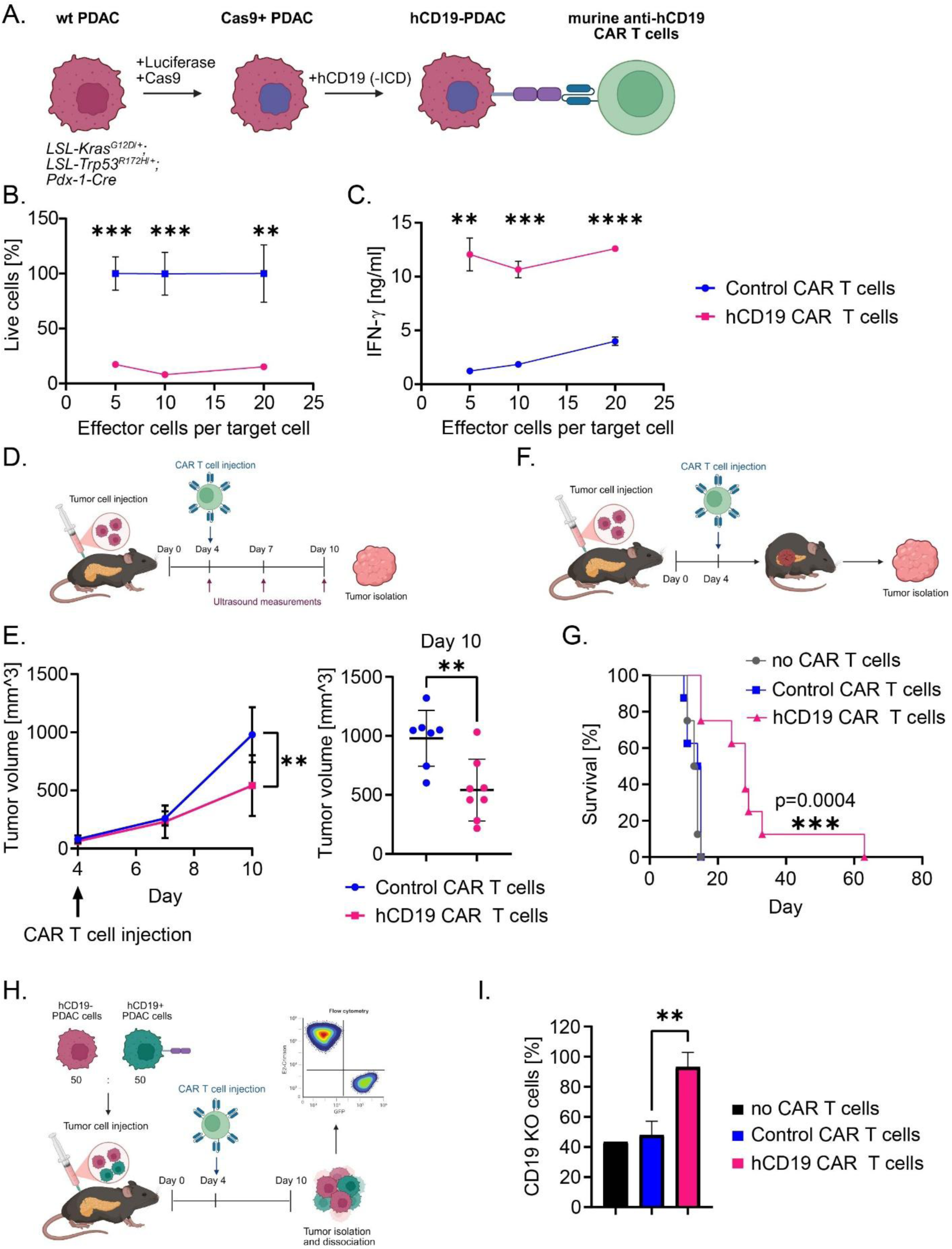
Development of a CRISPR-Cas9 knock-out screening-amenable PDAC model for studying CAR T cell therapy resistance. **A)** A KPC (*LSL-Kras^G12D/+^;LSL-Trp53^R172H/+^;Pdx1-Cre*) tumor-derived murine PDAC cell line was engineered to stably express *Firefly Luciferase*, *Sp. Cas9*, and hCD19, with a deletion of the intracellular domain (-ICD). The final cell line was established from a single cell cloned to ensure uniform Cas9 activity and CAR T cell antigen expression. **B)** A graph showing a CAR T cell cytotoxicity assay involving incubation of hCD19 CAR T cells or control (anti-EGFRvIII) CAR T cells with hCD19+ PDAC cells at multiple effector-to-target ratios (E:T ratios = CAR T cells:PDAC cells = 5:1, 10:1, 20:1) for 24 hours. All conditions were analyzed in triplicates and error bars represent standard deviation. Statistical analysis was conducted using unpaired t-test. **C)** A graph showing the level of IFN-γ release from hCD19 or control CAR T cells during the CAR T cell assay. All conditions were analyzed in triplicate and error bars represent standard deviation. Statistical analysis was conducted using unpaired t-test (ns > 0.05, * ≤ 0.05, ** p ≤ 0.01, *** p ≤ 0.001, **** p ≤ 0.0001). **D)** Experimental schematic: C57BL/6 mice received an orthotopic injection of 600,000 hCD19+ PDAC cells, followed by a subsequent intraperitoneal injection of freshly prepared hCD19 CAR T cells (n=8; 2.5 million) or control irrelevant CAR T cells (n=7) four days later. Tumor volume was measured every 3 days using ultrasound. **E)** (Left) A graph showing the tumor volume following CAR-T treatment. (Right) On day 10, hCD19+ PDAC tumors treated with hCD19 CAR-T cells are significantly (p=0.0059) smaller than control CAR T cell treated tumors. Error bars represent standard deviation. Statistical analysis was conducted using a Mann-Whitney test. **F)** Experimental schematic: C57BL/6 mice received an orthotopic injection of 600,000 hCD19+ PDAC cells, followed by a subsequent intraperitoneal injection of freshly prepared hCD19 CAR T cells (n=8; 2.5 million) or control irrelevant CAR T cells (n=8), or no CAR T cells four days later (n=8). Mice were monitored until a humane endpoint was reached and their survival was recorded. **G)** A Kaplan-Meier curve showing overall survival of mice bearing hCD19+ PDAC tumors treated with hCD19 CAR T cells, not treated or treated with control CAR T cells. CAR-T treatment significantly extends survival compared to treatment with control CAR T cells (Mantel-Cox test p=0.0004). **H)** Experimental schematic: PDAC cells with (GFP+) and without the hCD19 antigen (E2-Crimson+) were mixed at a 1:1 ratio. A total of 600,000 PDAC cells was orthotopically injected into C57BL/6 mice, followed by a subsequent intraperitoneal injection of freshly prepared hCD19 CAR T cells (n=5; 2.5 million) or control irrelevant CAR T cells (n=5) four days later. After ten days, PDAC tumors were isolated, dissociated into single cells, and analyzed by flow cytometry for GFP+ and E2-Crimson+ tumor cells. **I)** A bar graph showing that PDAC tumor cells without the CD19 antigen (E2-Crimson+) enrich over time in CAR T cell treated PDAC tumors. Error bars represent standard deviation. Statistical analysis was conducted using a Mann-Whitney test (p=0.0079).

To determine the feasibility of *in vivo* screening with hCD19+ PDAC using immunocompetent mouse cells, we examined the growth kinetics and tumor burdens of both Luc+ PDAC cells and hCD19+ PDAC cells. These cells exhibit similar *in vitro* proliferation rates (**Figure S1D**). Tumor growth kinetics were also similar between Luc+ and hCD19+ PDAC cells over a time course of 10 days, indicating that engineered PDAC cells exhibit no or very little immunogenicity *in vivo* (**Figure S1E**). Furthermore, we did not observe any differences in the weight of orthotopic PDAC tumors established from Luc+ or hCD19+ PDAC cells in either immunocompetent C57BL/6 mice or immunodeficient NOD-SCID/IL2Rg−/− (NSG) mice (**Figure S1F-G**). This lack of immunogenicity is in line with previously published work using a Cas9+ murine B cell acute lymphoblastic leukemia (B-ALL) model in C57BL/6 mice^23^.

Next, we investigated whether hCD19+ PDAC cells are susceptible to CAR T cell killing *in vitro*. We incubated 10,000 hCD19+ PDAC cells with increasing effector-to-target (E:T) ratios of murine CAR T cells targeting EGFRvIII, a control antigen not expressed by our cells, or the model antigen hCD19 for 24 hours (**Figure 1B**). We observed efficient killing of PDAC cells at all E:T ratios tested (5:1, 10:1 and 20:1), as well as robust IFN-γ release by hCD19 CAR T cells compared to control CAR T cells (**Figure 1C**).

To test whether hCD19+ PDAC cells could also be targeted by CAR T cells *in vivo*, we transplanted hCD19+ PDAC cells orthotopically into the pancreas of fully immunocompetent C57BL/6 mice followed by intraperitoneal injection of murine control or hCD19 CAR T cells four days later (**Figure 1D**). Tumor growth, monitored by ultrasound over the following ten days, was significantly delayed in mice treated with hCD19 CAR T cells compared to controls (**Figure 1E**), and this translated into a survival benefit (**Figure 1G**). Despite continued tumor enlargement during treatment, histological analysis revealed extensive necrosis, hemorrhage, and dense immune cell infiltration in CAR T cell–treated tumors (**Figure S1H**), consistent with robust intratumoral CAR T cell activity.

Having established that hCD19+ PDAC cells can be targeted by hCD19 CAR T cells *in vivo*, we then tested whether CAR T cell treatment can provide the selective pressure required to successfully identify mediators of response and resistance using CRISPR-Cas9 KO screening. A well-established mediator of resistance in patients treated with CAR T cell therapy is loss of the targeted tumor antigen. Therefore, antigen-negative tumor cells should enrich in a mixture with antigen-positive tumor cells when targeted with CAR T cells if the selective pressure is adequate in our model system. To test this, we mixed hCD19 antigen-positive (GFP+) PDAC cells with antigen-negative (E2-Crimson+) PDAC cells at a 1:1 ratio and treated them with control (EGFRvIII) or hCD19 CAR T cells at an E:T ratio of 10:1 for 24 hours *in vitro* (**Figure S2A**). Treatment with hCD19 CAR T cells, but not control CAR T cell treatment or no treatment, led to an increase in antigen-negative PDAC cells relative to antigen-positive PDAC cells (**Figure S2B**). Similarly, orthotopic injection of a 1:1 mixture of antigen-positive (GFP+) and antigen-negative (E2-Crimson+) PDAC cells followed by CAR T cell treatment with hCD19 or control CAR T cells lead to an enrichment of antigen-negative PDAC cells over a period of 10 days (**Figure 1H-I**). Together, these results confirm that CAR T cell treatment in our PDAC model can successfully enrich for an established resistance mechanism both *in vitro* and *in vivo*. Thus, this selective pressure should similarly enable the identification of novel mediators of CAR T cell therapy response and resistance using CRISPR-Cas9 KO screening.

For successful *in vivo* CRISPR-Cas9 KO screening, it is essential to maintain an adequate representation of single guide RNAs (sgRNAs) throughout the experiment^37^. The size limit of the sgRNA library that can be screened robustly *in vivo* is influenced by the tumor type, the number of tumor cells that can be safely injected, and the engraftment efficiency of tumor cells in the chosen location. To empirically determine the maximum library size that can be screened in the hCD19+ PDAC model, we generated individual hCD19+ PDAC cell lines expressing either GFP+ or mCherry+ to perform limiting GFP-dilution assays *in vivo*. To do so, we established five populations of hCD19+ PDAC cells containing decreasing fractions of GFP+ cells (50%, 1%, 0.1%, 0.05%, or 0.025%) mixed with mCherry+ cells, and orthotopically transplanted them into C57BL/6 mice (**Figure S2C**). Each GFP percentage corresponds to the maximal empirical library size that could be used if the corresponding GFP percentage is recovered at the experimental endpoint (**Figure S2D**). Ten days after transplantation, PDAC tumors were isolated and analyzed by flow cytometry for GFP+ PDAC cells. We then compared the percentage of GFP+ cells recovered at the end-point (“output”) to the starting percentage of GFP+ cells prior to orthotopic transplantation (“input”) (**Figure S2E**). In all experimental groups, the empirical output to input ratio was ∼1, indicating robust *in vivo* maintenance of GFP+ PDAC cells throughout the entire experiment. These results indicate that our hCD19+ PDAC model could adequately represent a library of ∼4000 sgRNAs at 150x coverage *in vivo*. Taken together, we have established an immunocompetent, orthotopic PDAC model amenable to CRISPR-Cas9 KO screening in the context of CAR T cell therapy.

### *In vivo* CRISPR-Cas9 screens reveals modulators of CAR T cell response in PDAC

Next, we performed iterative KO screens using the hCD19+ PDAC model system to uncover genetic modulators of CAR T cell response in pancreatic cancer. For this purpose, we made use of the modular SKY CRISPR library that is divided into non-redundant pools that can be individually screened or combined into larger libraries^23^. This genome-wide library contains four sgRNAs per protein-coding murine genes, divided into 48 pools. Using their initial Kyoto Encyclopedia of Genes and Genomes (KEGG) term in a non-redundant manner, 36% of protein-coding genes could be classified into a KEGG pathway, comprising the first 14 pools and part of pool 15 covering all KEGG pathways, while all other genes were randomly distributed among the remaining pools. Each pool also contained control sgRNAs targeting intergenic regions, olfactory genes, and murine essential genes^38^, and the first pool additionally contained sgRNAs targeting hCD19.

We performed a total of eight independent KO screens covering the first 16 pools of the SKY library (**Table S1**). Guided by our limiting dilution experiments (**Figure S2E-F**), we decided to combine and screen two library pools (∼3800 total sgRNAs) in individual mice. Each screen was performed by transplanting 600,000 sgRNA-expressing hCD19+ PDAC cells orthotopically into the pancreas of C57BL/6 mice followed by treatment with murine hCD19 CAR T cells or control CAR T cells targeting EGFRvIII four days later (**Figure 2A**). Each batch of freshly produced murine CAR T cells was functionally validated for antigen-dependent on-target killing efficiency and IFN-γ release *in vitro* before use *in vivo* (**Figure S3A-C**). Tumors were isolated 10-25 days after initial transplantation, when mice showed advanced tumor burden (**Figure S3D-K**), dissociated into single cell suspensions, and subjected to high-throughput sequencing to quantify sgRNA representation. sgRNA coverage was well maintained across all experimental arms compared to pre-screen input samples (**Figure S4A-B**). To further assess screen quality, we confirmed that many sgRNAs targeting essential genes were significantly depleted in both control and hCD19 CAR T cell-treated groups, whereas those targeting non-essential genes remained unchanged (**Figure S5A**).

**Figure 2:**
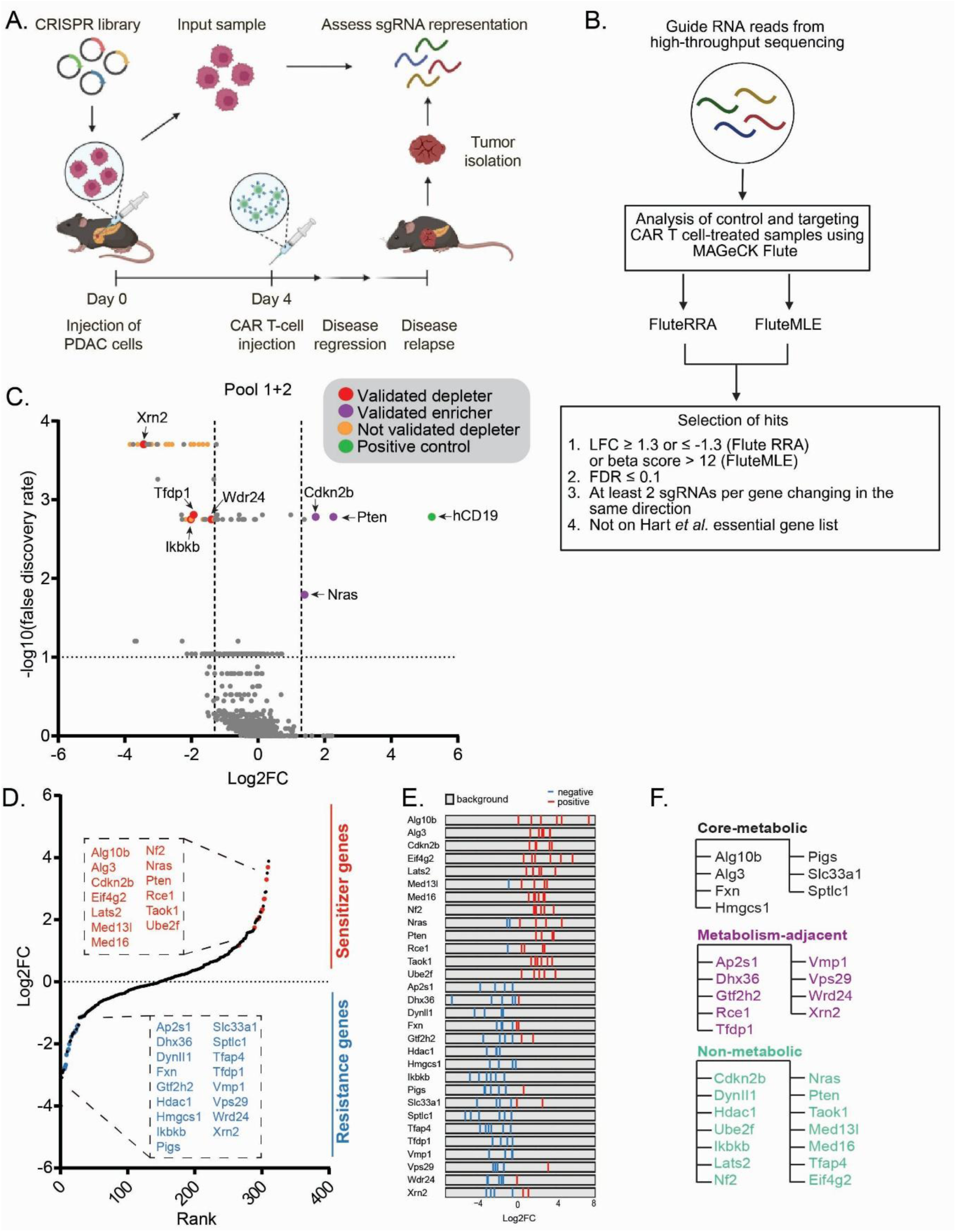
*In vivo* CRISPR-Cas9 KO screening reveals modulators of response to CAR T cell therapy in PDAC. **A)** Experimental schematic: murine PDAC cells containing CRISPR library pools were injected orthotopically into C57BL/6 mice, followed by intraperitoneal injection of hCD19 CAR T cells or control CAR T cells (n=8 mice per two pools, per CAR T cell treatment) four days later. Mice were monitored, and tumors were harvested when a humane endpoint was reached. PDAC tumors were dissociated into single cell suspensions, genomic DNA was isolated and guide RNA representation was determined by next-generation sequencing (NGS). Input samples were harvested of PDAC cells containing the CRISPR libraries on the day of orthotopic injection and also subjected to NGS. In total, eight independent screens were performed, covering more than 6000 genes associated with KEGG pathways. **B)** A schematic of our analysis pipeline: guide RNA reads were analyzed using MAGeCK Flute algorithms. Both the FluteRRA and FluteMLE algorithms were used to determine significantly changed guide RNAs between control CAR T cell treated and hCD19 CAR T cell treated PDAC tumor cells. Genes were considered a “hit” if they fulfilled four criteria: (1) an LFC of greater than 1.3 or smaller than -1.3 (FluteRRA) or a beta score higher than 12 (FluteMLE), (2) an FDR equal to or smaller than 0.1, (3) at least two out of four guide RNAs changing in the same direction, and (4) are not considered essential genes (Hart et al.). **C)** A volcano plot showing the results of the first knock-out screen with CRISPR-Cas9 library pools 1 and 2. Library pool 1 contains guide RNAs targeting the CAR T cell antigen hCD19 as a positive control. “Validated depleter/enricher” refers to genes that were validated in a secondary CRISPR-Cas9 KO screen. “Not validated depleter” refers to genes that were not confirmed as hits in a secondary CRISPR-Cas9 KO screen. **D**) A rank plot (left) of all genes re-screened in a validation CRISPR-Cas9 library under the same conditions described in A. The validation library covers all enrichers and depleters identified in the eight primary CRISPR-Cas9 KO screens and contains six guide RNAs per gene. All genes that validated in this secondary screen are highlighted and listed. In total, 13 genes that promote CAR T cell response and 17 genes that promote CAR T cell resistance were identified. **E)** Log2 fold changes of individual guide RNAs for each resistance and sensitizer gene identified in the validation screen. Guide RNA enrichment is shown in red and guide RNA depletion in blue. **F)** Classification of modulators of CAR T cell therapy response into core-metabolic, metabolism-adjacent or non-metabolic genes using KEGG, Reactome, and GO annotations.

Next, we used the Model-based Analysis of Genome-wide CRISPR-Cas9 Knockout (MAGeCK) FluteRRA and FluteMLE algorithms to determine significantly enriched or depleted sgRNAs between control CAR T cell-treated and hCD19 CAR T cell-treated PDAC tumor cells (**Figure 2B**)^39^. Enriched sgRNAs are predicted to target genes whose inactivation mediates resistance to CAR T cell therapy, while depleted sgRNAs target genes whose loss sensitizes PDAC cells to CAR T cell therapy. Genes were considered a “hit” if they fulfilled four rigorous criteria: (1) a log-fold change (LFC) of greater than 1.3 or smaller than -1.3 (as determined by FluteRRA) or a beta score higher than 12 (as determined by FluteMLE), (2) a False Discover Rate (FDR) equal to or smaller than 0.1, (3) at least two out of four sgRNAs changing in the same direction, and (4) not part of the essential gene catalog defined by Hart *et al*^38^. Importantly, sgRNAs targeting the hCD19 CAR T cell antigen showed the strongest enrichment of all sgRNAs in the first screen (**Figure 2C**). These results confirm that adequate selective pressure was generated with CAR T cell therapy in our hCD19+ PDAC model, allowing us to identify genetic modulators of CAR T cell response *in vivo*. Across all eight KO screens, a total of 244 genes fulfilled our “hit” selection criteria and altered the representation of hCD19+ PDAC cells following CAR T cell therapy (**Figure S6, Table S2**). Of these 244 potential CAR T cell therapy modulators, 30 genes were classified as “sensitizers” and the remaining 204 genes were classified as “resistance” genes.

A caveat of our pooled screening approach is that it does not enable a ranked comparison between sgRNAs across separate screens. Therefore, we performed a validation screen using a new CRISPR KO library containing six sgRNAs targeting each gene previously identified as a “hit” (**Table S2**), along with 90 non-targeting (scrambled), 213 intergenic, and 91 olfactory gene-targeting sgRNAs that exhibited neutral behavior (no enrichment or depletion) in our initial screens. We performed this validation screen in an identical fashion to the primary KO screens. CAR T cell functionality was confirmed *in vitro* by efficient on-target killing and IFN-γ release (**Figure S7A-C**). Tumors were isolated when mice showed advanced tumor burden, 15-32 days after initial injection (**Figure S7D**), and sgRNA representation was quantified using high-throughput sequencing. Compared to pre-screen input samples, sgRNA coverage was generally maintained, although some variation was observed, especially in tumors from long-term survivors treated with hCD19 CAR T cells (**Figure S7E**). As in our primary screens, sgRNAs targeting known essential genes have large negative fold changes relative to non-essential genes in both the control and hCD19 CAR T cell treated groups compared to their respective pre-screen input samples, whereas sgRNAs targeting intergenic regions or non-essential genes were unchanged, confirming screen robustness (**Figure S7F**).

*Bona fide* modulators of CAR T cell therapy were determined by assessing the FDR values, LFCs, and both intra-screen (within validation screen) and inter-screen (compared to primary screens) consistency of sgRNA behavior (**Figure S7G**). Of the 244 genes that we screened, 30 genes validated as robust modulators of CAR T cell therapy (**Figure 2D**) 13 of these genes sensitized tumor cells to CAR T cell therapy as indicated by consistent enrichment of sgRNAs targeting these genes (**Figure 2E**). This group consisted of known tumor suppressor genes (e.g. *Pten*, *Cdkn2b, Eif4g2*)^40–42^, and genes regulating cell proliferation (e.g. Hippo pathway genes *Nf2, Taok1, Lats2*)^43^. Importantly, loss of *Pten* has previously been shown to mediate resistance to immune checkpoint blockade (ICB) therapy in multiple tumor types^44–47^. The remaining 17 genes were identified as mediators of resistance to CAR T cell therapy, as indicated by consistent depletion of sgRNAs targeting those genes (**Figure 2E**). This set included mediators of autophagy (*Vmp1, Wrd24*) and intracellular trafficking (*Vps29*) processes previously shown to mediate immune evasion from T cells^48^. We also identified known regulators of clathrin-mediated endocytosis (*Aps2s1, DynII1*) as resistance genes to CAR T cell therapy^49^. Surprisingly, multiple genes identified in our screen were core-metabolic genes (**Figure 2F**), encoding enzymes or transporters in canonical metabolic pathways defined by KEGG, Reactome, and GO annotations (*Hmgcs1, Sptlc1, Slc33a1, Fxn, Pigs, Alg10a, and Alg3*)^50–53^. Additionally, nine other genes can be classified as metabolism-adjacent due to their roles in nutrient sensing, autophagy, and metabolic stress adaptation (**Table S3**). Currently, no tumor-intrinsic alterations in metabolic genes have been described to influence CAR T cell response, highlighting metabolic stress regulation as a previously unrecognized determinant of PDAC susceptibility to CAR T cell therapy.

### Single cell RNA sequencing of CAR T cell treated PDAC tumors reveals changes in cellular redox states driven by Nrf2

The identification of metabolic and stress-response genes as modulators of CAR T cell response in our CRISPR screen raised the possibility that CAR T cell therapy imposes selective pressure on tumor-intrinsic metabolic states in PDAC. To test this using an orthogonal methodology, we performed scRNA seq of orthotopic hCD19+ PDAC tumors treated with control (EGFRvIII) or hCD19 CAR T cells. For this purpose, we injected hCD19+ PDAC cells orthotopically into C57BL/6 mice and treated them with freshly produced murine CAR T cells (**Figure 3A, Figure S8A-C**). Tumors were isolated after 10 days for control CAR T cell-treated mice and after 15 days for hCD19 CAR T cell-treated mice, at which stage all PDAC tumors had reached a similar weight (**Figure S8D**). Following tumor dissociation into single cells, live cell mixtures were subjected to scRNA seq. The resulting data were filtered and 20,516 cells were portioned into clusters of similar cells (**Figure S8E; see Methods**). We identified tumor cells by hCD19 and Cas9 expression, and annotated stromal and immune cell clusters using well-established genes and gene signatures, such as *Cd19* for B-cells and *Cd68* for macrophages (**Figure S8F and Figure S9A**). In total, we identified 12 transcriptionally distinct clusters consisting of PDAC cells, cells of the innate and adaptive immune system, including CAR T cells (*Cd8a* and GFP expression), endothelial cells, pancreatic stellate cells and erythroid cells (**Figure S9B**).

**Figure 3:**
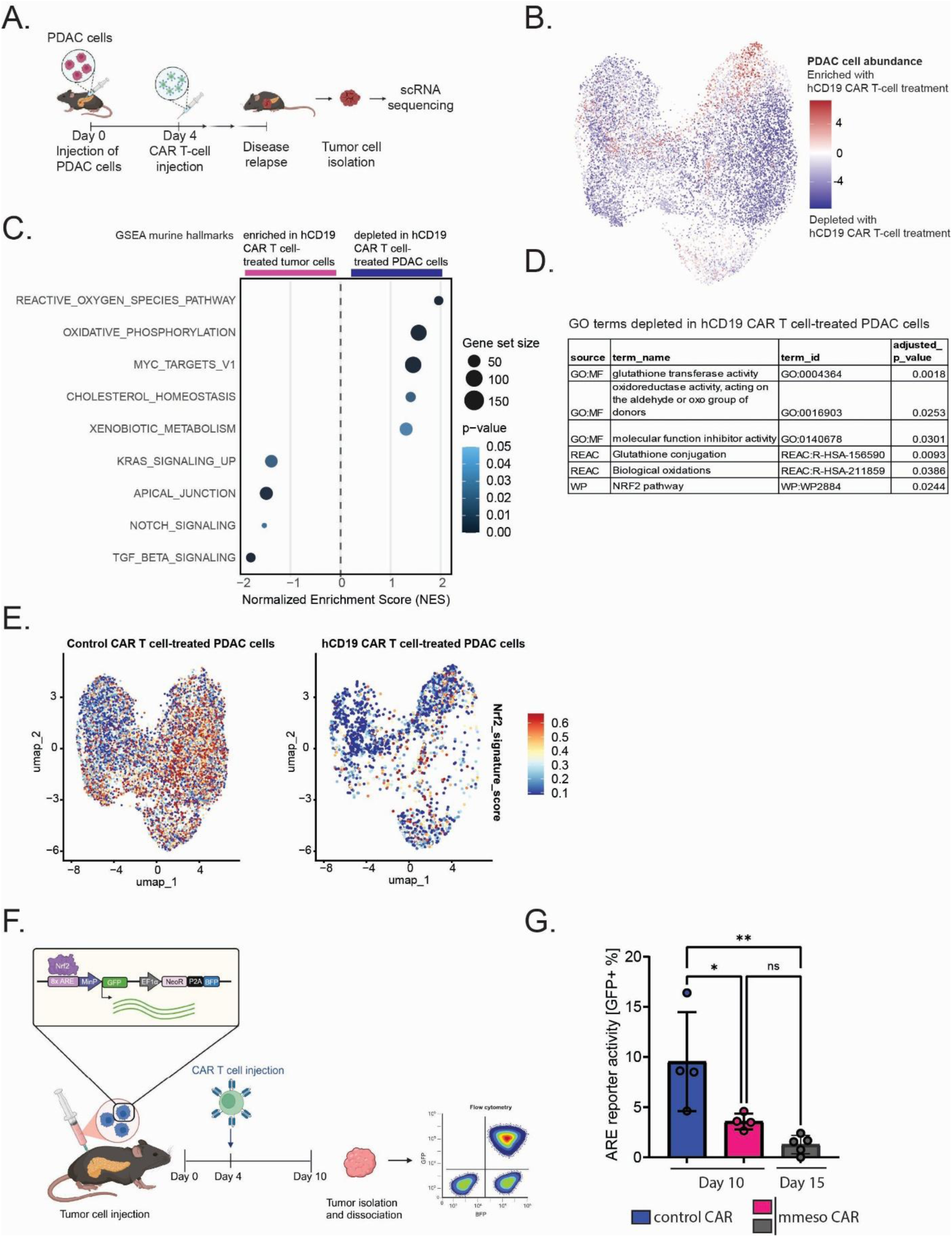
Single cell RNA sequencing of CAR T cell treated PDAC tumors reveals changes in cellular redox states. **A)** Experimental schematic: C57BL/6 mice received an orthotopic injection of 600,000 murine PDAC cells, followed by a subsequent intraperitoneal injection of freshly prepared hCD19 CAR T cells (n=4; 2.5 million) or control irrelevant CAR T cells (n=4) four days later. PDAC tumors were harvested once tumors reached approx. 1500 mm^3^ (day 10 for control CAR T cell treated tumor; day 15 for hCD19 CAR T cell treated tumors). Tumors were dissociated into single cell suspensions and chromium single cell RNA sequencing (10x genomics) was performed. **B)** A UMAP of PDAC cells from tumors treated with hCD19 CAR T cells or control CAR T cells. The differential abundance of tumor cells between treatment conditions is depicted by color in the UMAP. Tumor cells enriching with hCD19 CAR T cell treatment compared to control CAR T cell treatment are colored in red, and tumor cells depleting following hCD19 CAR T cell treatment are colored in blue. Differentially expressed genes were identified between PDAC cells of these two experimental conditions and used for pathway analyses. **C)** Dot plot showing Gene Set Enrichment Analysis (GSEA) results of murine hallmarks in PDAC cells in hCD19 CAR T cell treated tumors compared to control CAR T cell treated tumors. **D)** Gene ontology (GO) analysis of top 100 depleted genes in hCD19 CAR T cell treated tumors compared to control CAR T cell treated tumors. **E)** A UMAP showing the expression of a Nrf2-responsive gene signature (murine orthologs of the human NRF2 Wikipathway) in PDAC cells from tumors treated with hCD19 CAR T cells or control CAR T cells. Nrf2-responsive gene expression is lower in PDAC cells treated with hCD19 CAR T cells compared to control CAR T cell treated PDAC cells. **F)** Experimental schematic: C57BL/6 mice received an orthotopic injection of 600,000 murine PDAC cells expressing a *Nrf2* 8xARE-GFP reporter, followed by a subsequent intraperitoneal injection of freshly prepared mmeso CAR T cells (5 million) or control irrelevant CAR T cells four days later (n=5 per experimental group). Binding of Nrf2 to the ARE sites of the reporter leads to the expression of GFP. Tumors were isolated on day 10 or day 15 and dissociated into single cell suspensions, and Nrf2 activity (GFP) in PDAC cells (BFP+) was analyzed using a flow cytometer. **G)** A bar graph showing Nrf2 activity (as measured by GFP expression) in PDAC cell isolated from tumors treated with mmeso CAR T cells compared to control CAR T cell treated tumors. Statistical analysis was conducted with Ordinary One-Way ANOVA test (ns > 0.05, * ≤ 0.05, ** p ≤ 0.01, *** p ≤ 0.001, **** p ≤ 0.0001).

To identify differentially expressed genes between PDAC cells treated with control CAR T cells compared to hCD19 CAR T cells, we isolated the 13,410 PDAC cells in our dataset and performed unsupervised clustering and differential gene expression analysis. Furthermore, we identified differences in the abundance of tumor cells in different regions of the 2-dimensional uniform manifold approximation and clustering (UMAP) plots between control and hCD19 CAR T cell treatment, showing CAR T cell treatment specific enrichment and depletion of certain tumor cell clusters (**Figure 3B**). To identify biological processes and molecular functions impacted in PDAC cells upon treatment with CAR T cells, we performed Gene Set Enrichment Analysis (GSEA) on murine hallmark genes. This analysis showed the depletion of reactive oxygen species, oxidative phosphorylation, *Myc* targets, cholesterol homeostasis, and xenobiotic metabolism pathways in PDAC cells following hCD19 CAR T cell treatment, while signaling pathways such as *Tgf-β, Notch,* apical junction, and *Kras* signaling were enriched (**Figure 3C**). In parallel, we performed gene ontology (GO) analysis using gProfiler on the top 100 differentially expressed genes revealed downregulation of pathways related to oxidative stress in hCD19 CAR T cell-treated PDAC cells, including glutathione metabolism, oxidoreductase activity, biological oxidation, and the Nrf2 (nuclear factor, erythroid 1-like2) pathway (**Figure 3D**). Conversely, pathways regulating the interaction of tumor cells with the extracellular matrix were upregulated in hCD19 CAR T cell treated PDAC cells (**Figure S9C**). Together, these results indicate that PDAC cells treated with hCD19 CAR T cells exhibit substantial molecular changes in oxidative stress pathways, including the Nrf2 antioxidant response.

### Nrf2-activated PDAC cells are eliminated by CAR T cell therapy

The Nrf2-antioxidant response element (ARE) signaling pathway plays a key role in the cellular defense against oxidative or electrophilic stress^54^. The cytosolic protein Kelch-like ECH Associated-Protein 1 (Keap1) acts as the central negative regulator of this pathway by targeting Nrf2 for proteasomal degradation^55^. Intriguingly, mutations in *Keap1* that lead to Nrf2 hyperactivation have been implicated in mediating resistance to ICB therapy in lung adenocarcinoma (LUAD)^56^. This resistance can be overcome by inhibition of glutaminase, a key enzyme that produces glutamate which is required for the glutathione redox system^57^. To further analyze the connection between the Nrf2 pathway and CAR T cell therapy, we investigated the expression of a gene signature of murine Nrf2-responsive genes in PDAC cells following hCD19 CAR T cell treatment (**Figure 3E, Table S4**). Consistent with our previous results, expression of murine Nrf2-responsive genes was significantly reduced in PDAC cells that survived hCD19 CAR T cell treatment compared to control CAR T cell treatment. Thus, we hypothesized that PDAC cells with Nrf2 pathway activation may show increased sensitivity to CAR T cell treatment. To test this, we leveraged a previously described *Nrf2* 8xARE-GFP reporter for which GFP expression serves as a readout of *Nrf2* transcriptional activity^58^. Following transduction of WT PDAC cells with the *Nrf2* 8xARE-GFP reporter (*Nrf2* reporter), we validated reporter activity by treating these cells with tert-butylhydroquinone (tBHQ), an established *Nrf2* activator^59^. We observed a robust, dose-dependent induction of GFP expression *in vitro*, thereby validating the assay (**Figure S10A-B**). Next, we orthotopically transplanted *Nrf2* reporter PDAC cells into C57BL/6 mice and treated them with either control (EGFRvIII) or murine mesothelin (mmeso) CAR T cells (**Figure 3F**)^60^. Ten days later, we isolated the tumors and analyzed *Nrf2* reporter activity via GFP expression in PDAC cells. Strikingly, *Nrf2* reporter activity was significantly lower in PDAC cells isolated from tumors treated with mmeso CAR T cells compared to those isolated from tumors treated with control CAR T cells at the same time point (**Figure 3G**). The PDAC tumors treated with mmeso CAR T cells were significantly smaller than control CAR T cell treated tumors on the same day (**Figure S10C**). To match these data to the time points of our previous scRNA seq experiment, we isolated PDAC tumors treated with mmeso CAR T cells after 15 days as well, at which point the tumors had reached a similar size as to control CAR T cell treated tumors on day 10 (**Figure S10C**). Interestingly, we observed an even further decrease of Nrf2 reporter activity in mmeso CAR T cell treated tumors on day 15 (**Figure 3G**). These results indicate that PDAC cells with high *Nrf2* pathway activity may be hypersensitive to, and selectively eliminated by, CAR T cell therapy *in vivo*. This stands in contrast to ICB in which Nrf2-hyperactivation in cancer cells is associated with therapy resistance^61^.

### *Slc33a1* loss and Nrf2-hyperactivation are synthetically lethal in PDAC

Our CRISPR-Cas9 KO screening and single cell transcriptome analysis of PDAC cells treated with CAR T cell therapy converge on metabolic genes as critical modulators of CAR T cell therapy response. Of particular note, *Slc33a1*, a mediator of resistance to CAR T cell therapy in our screen, has recently been shown to be a key mediator of tolerance to Nrf2-hyperactivation states. Specifically, *Slc33a1* inactivation was shown to be synthetically lethal with loss-of-function mutation of *Keap1* in LUAD^62^. Additionally, Nrf2-hyperactivation is also observed in PDAC tumors^63^. Therefore, we wanted to confirm that Slc33a1 was similarly linked to the tolerance of Nrf2-hyperactive states in PDAC.

To confirm the synthetic lethal interaction between *Slc33a1* KO and *Keap1* KO in hCD19+ PDAC cells, we performed *in vitro* competition assays with hCD19+ PDAC cells containing a control (*Olfr402*) KO, *Slc33a1* KO, or *Slc33a1* re-expression in combination with *Keap1* KO. PDAC cells with *Slc33a1*/*Keap1* double-KO depleted over time compared to PDAC cells with single *Slc33a1* KO whereas *Olfr402/Keap1* double-KO cells did not deplete in comparison to *Olfr402* KO PDAC cells (**Figure S10D**). At the experimental endpoint, the remaining PDAC cells in the *Keap1/Slc33a1* double-KO population were selected against *Keap1* loss (**Figure S10E**). These results demonstrate that the *Keap1*/*Slc33a1* synthetic lethal interaction is conserved across different types of tumor cells.

### Loss of *Slc33a1* sensitizes PDAC tumors to CAR T cell therapy *in vivo*

*Slc33a1* encodes for the solute carrier family member 1 protein, an acetyl-coenzyme A transporter located in the endoplasmic reticulum (ER) membrane^64^. Its function is essential for the lysine acetylation of proteins in the ER, and both loss-of-function or overactivation of *Slc33a1* have been implicated in various diseases^65,66^. In pancreatic acinar cells, *Slc33a1* KO has been shown to result in ER stress, fibrosis, and inflammation^67^.

To validate *Slc33a1* as a resistance mediator to CAR T cell therapy, we injected hCD19+ PDAC cells with *Slc33a1* KO (**Figure 4A**) or control (*Olfr402*) KO into C57BL/6 mice and treated them with murine control (EGFRvIII) or hCD19 CAR T cells (**Figure 4B**). We observed a statistically significant extension of survival in mice injected with *Slc33a1* KO hCD19+ PDAC cells compared to control KO PDAC cells treated with hCD19 CAR T cells (**Figure 4C**). Mice injected with *Slc33a1* KO hCD19+ PDAC cells treated with hCD19 CAR T cells ultimately succumbed to disease; however, end-stage *Slc33a1* KO tumors were significantly larger compared to all other experimental groups and showed extensive immune cell infiltration, hemorrhage, and necrosis (**Figure 4D-E**). These features are suggestive of pseudo progression, a clinically observed phenomenon during immunotherapy in which immune-mediated inflammation and cell death lead to apparent increases in tumor volume despite effective antitumor responses^68–70^. In this context, the absence of tumor regression may not reflect therapeutic failure, but rather that tumors reach a size incompatible with survival before meaningful shrinkage can occur. While we cannot definitively distinguish pseudo progression from true progression in this model, the observed histological changes and survival benefit support this as a potential explanation.

**Figure 4:**
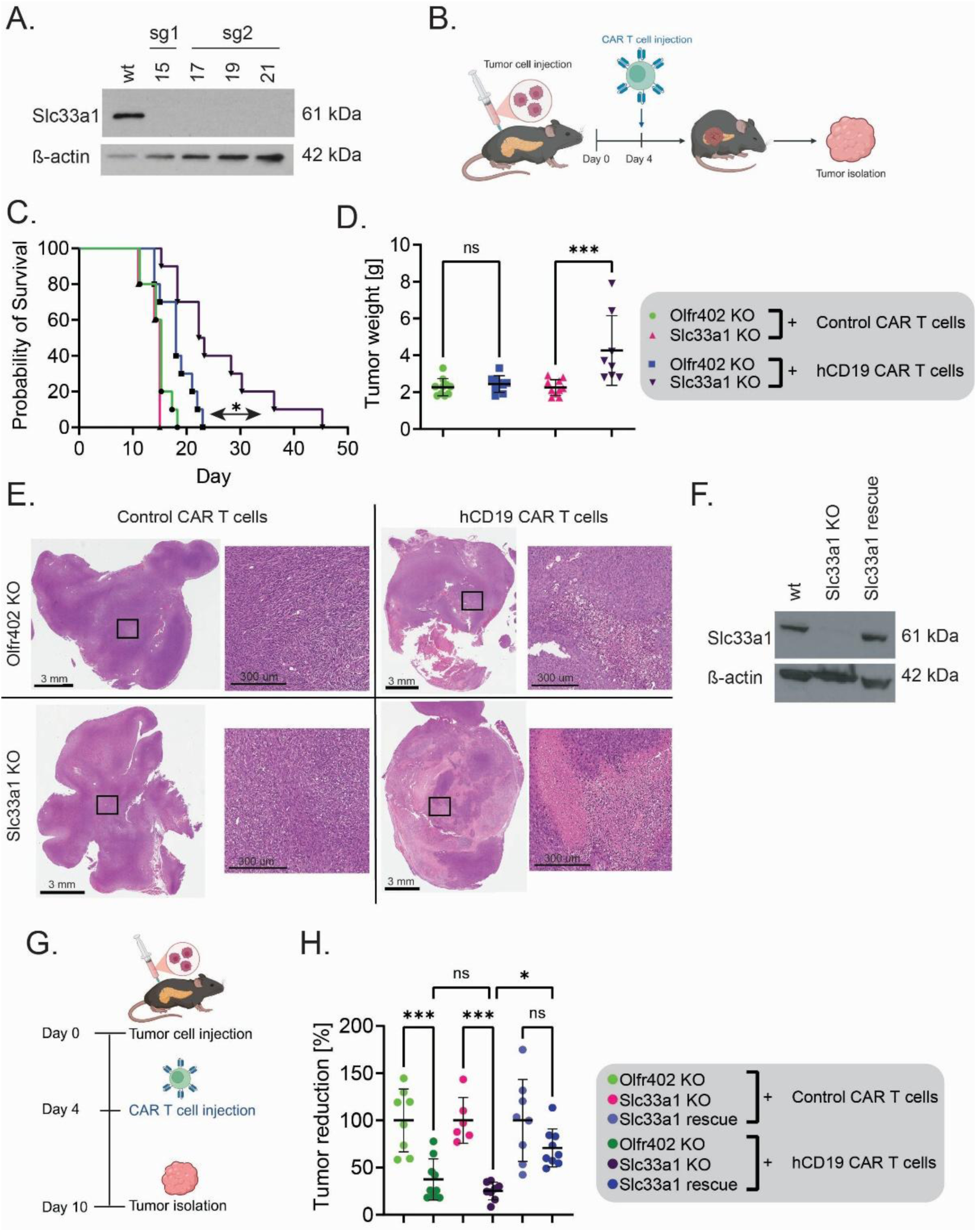
Loss of *Slc33a1* sensitizes to CAR T cell therapy *in vivo*. **A)** Western blot analysis showing knock-out of *Slc33a1* in multiple hCD19+ PDAC single cell clones using two different sgRNAs. **B)** Experimental schematic: C57BL/6 mice received an orthotopic injection of 600,000 murine PDAC cells with a control KO (*Olfr402*) or *Slc33a1* KO, followed by a subsequent intraperitoneal injection of freshly prepared hCD19 CAR T cells (5 million) or control (EGFRvIII) irrelevant CAR T cells four days later (n=10 per experimental group). Mice were monitored until a humane endpoint was reached and their survival was recorded. **C)** A Kaplan-Meier curve showing survival in mice bearing *Slc33a1* KO or control KO tumors treated with control or anti-CD19 CAR-T cells. *Slc33a1* KO in PDAC cells extends survival in C57BL/6 mice treated with hCD19 CAR T cell. Mantel-Cox test p=0.0123. **D)** A graph showing PDAC tumor weight at morbidity in mice treated with hCD19 CAR T cells or control CAR T cells. PDAC tumors containing *Slc33a1* KO PDAC cells treated with hCD19 CAR T cells are significantly larger p=0.0006. Statistical analysis was conducted with Ordinary One-Way ANOVA test (ns > 0.05, * ≤ 0.05, ** p ≤ 0.01, *** p ≤ 0.001, **** p ≤ 0.0001). **E)** H&E-stained sections of tumors containing PDAC cells with control (*Olfr402*) KO or *Slc33a1* KO treated with hCD19 CAR T cells or control CAR T cells. The left side of each panel shows the whole tumor cross-section and the right side zooms into a specified tumor area (black square) demarcated on the left side. PDAC tumors containing *Slc33a1* KO PDAC cells treated with hCD19 CAR T cells show large necrotic areas. **F)** Western blot analysis showing reconstitution (rescue) of *Slc33a1* in *Slc33a1* KO cells (sg1 cl15) compared to wt hCD19+ PDAC cells. **G)** Experimental schematic: C57BL/6 mice received an orthotopic injection of 600,000 murine PDAC cells with a control KO (*Olfr402*), *Slc33a1* KO, or *Slc33a1* rescue followed by a subsequent intraperitoneal injection of freshly prepared hCD19 CAR T cells (5 million) or control irrelevant CAR T cells four days later (n=6-8 per experimental group). Tumors were harvested on day 10 post-PDAC cell injection. **H)** Relative tumor reduction for hCD19 CAR T cell-treated PDAC tumors relative to control CAR T cell treated tumors. Statistical analysis was conducted with Ordinary One-Way ANOVA test and only significant differences are annotated (* ≤ 0.05, **** p ≤ 0.0001).

Next, we tested whether *Slc33a1* loss in tumor cells causes any changes in the tumor microenvironment that may contribute to their increased susceptibility to CAR T cell therapy. To control for potential clonal effects occurring in *Slc33a1* KO cells, we expressed a Cas9-resistant version of *Slc33a1* in *Slc33a1* KO cells to create *Slc33a1* rescue cells (**Figure 4F**). Then we injected hCD19+ PDAC cells with control (*Olfr402*) KO, *Slc33a1* KO, or *Slc33a1* rescue orthotopically into C57BL/6 mice, treated them with murine control (EGFRvIII) or hCD19 CAR T cells, and isolated the tumors after 10 days (**Figure 4G**). On-target (hCD19) CAR T cell therapy consistently reduced the tumor weight of control KO and *Slc33a1* KO PDAC tumors, but not of *Slc33a1* rescue tumors (**Figure 4H, Figure S11A**). Here, *Slc33a1* rescue tumors were the most resistant to hCD19 CAR T cell therapy.

Next, we dissociated these tumors and analyzed their tumor cell and immune cell content by flow cytometry. No differences in tumor cell content or CD45+ immune cells, including T cell subsets, B cells, NK cells and myeloid cell populations, were observed between control KO, *Slc33a1* KO or *Slc33a1* rescue tumors treated with control CAR T cells (**Figure S11B-L**). Similarly, there were no significant changes in immune cell populations between control KO tumors and *Slc33a1* KO tumors treated with hCD19 CAR T cells. The only changes in immune cell subsets we observed were caused by hCD19 CAR T cell treatment and were independent of, *Slc33a1* status, suggesting that *Slc33a1* loss does not alter the immune tumor microenvironment.

Given that the enhanced sensitivity of *Slc33a1*-deficient cells is not caused by the immune tumor microenvironment, we wanted to explore cell-intrinsic transcriptome changes caused by *Slc33a1* KO in tumor cells *in vivo*. Therefore, we injected hCD19+ PDAC cells with control (*Olfr402*) KO or *Slc33a1* KO orthotopically into C57BL/6 mice and treated them with murine control (EGFRvIII) or hCD19 CAR T cells. After 12 days, we harvested the tumors, isolated E2-Crimson+ tumor cells by fluorescence-activated cell sorting, and performed bulk transcriptome profiling. Differential gene expression analysis between control KO and *Slc33a1* KO tumor cells followed by GSEA revealed a depletion in inflammatory response signaling, such as IFN-γ, IFN-α, and IL6-JAK-STAT signaling, but an enrichment in the UPR in PDAC cells with *Slc33a1* loss (**Figure S12A-C**). This is in line with previous studies showing that *Slc33a1* loss leads to ER stress and the induction of the UPR^62,67^.

### Nrf2 hyperactivation sensitizes PDAC tumors to CAR T cell therapy *in vivo*

Given that (1) Nrf2-high PDAC cells may be selectively sensitive to CAR T cells, (2) *Keap1* is a negative regulator of *Nrf2* response, and (3) there is a conserved genetic link between *Slc33a1* and *Keap1*, we tested the mechanistic hypothesis that *Keap1* loss alone could sensitize PDAC cells to CAR T cell therapy. To test this, we orthotopically transplanted hCD19+ PDAC cells with control (*Olfr402*) or previously generated *Keap1* KO (sg2) into C57BL/6 mice and treated them with control (EGFRvIII) or hCD19 CAR T cells (**Figure 5A-B**). Consistent with our hypothesis, mice harboring *Keap1* KO PDAC tumors treated with hCD19 CAR T cells showed a significant survival advantage compared to mice with control PDAC tumors treated with hCD19 CAR T cells (p=0.0086; **Figure 5C**). Of note, while *Keap1*-targeting sgRNAs were not present in our initial screening library, the orthogonal use of scRNA seq and the genetic interaction between *Keap1* and *Slc33a1* allowed us to identify a human cancer gene and a mechanistic cellular redox state that modulates response to CAR T cell therapy.

**Figure 5:**
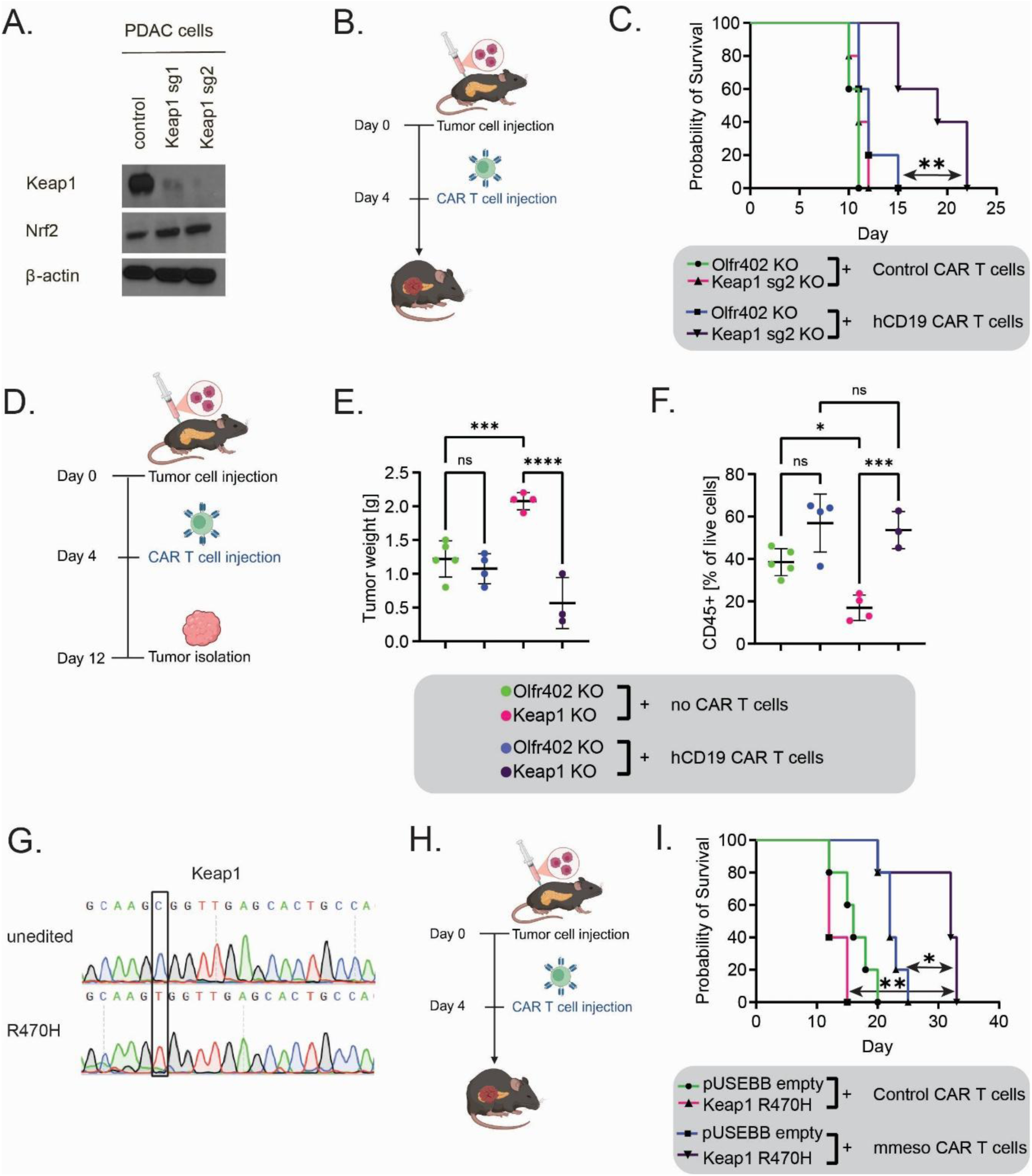
Nrf2 hyperactivation sensitizes to CAR T cell therapy *in vivo*. **A)** Western blot analysis showing *Keap1* levels using two separate sgRNAs in a bulk population of murine PDAC cells. **B)** Experimental schematic: C57BL/6 mice received an orthotopic injection of 600,000 murine PDAC cells with a control (*Olfr402*) KO or *Keap1* KO, followed by a subsequent intraperitoneal injection of freshly prepared hCD19 CAR T cells (5 million) or control (EGFRvIII) CAR T cells four days later (n=5 per experimental group). Mice were monitored until a humane endpoint was reached and their survival was recorded. **C)** A Kaplan-Meier curve showing survival of mice bearing Keap1 KO or control PDAC tumors treated with control or hCD19 CAR-T cells. Survival of mice injected with murine PDAC cells with control (*Olfr402*) KO or *Keap1* KO (sg2) shows that *Keap1* loss significantly extends survival in combination with hCD19 CAR T cell treatment (Mantel-Cox test p=0.0086). **D)** Experimental schematic: C57BL/6 mice received an orthotopic injection of 600,000 murine PDAC cells with a control KO (*Olfr402*), or *Keap1* KO followed by a subsequent intraperitoneal injection of freshly prepared hCD19 CAR T cells (2 million) or no CAR T cells (saline) four days later (n=3-5 per experimental group). Tumors were harvested on day 12 post-PDAC cell injection. **E)** Tumor weight of previously harvested PDAC tumors. Statistical analysis was conducted with Ordinary One-Way ANOVA test (ns > 0.05, * ≤ 0.05, ** p ≤ 0.01, *** p ≤ 0.001, **** p ≤ 0.0001). **F)** Flow cytometric analysis of CD45+ immune cells in *Keap1* KO and control (*Olfr402*) KO tumors treated with targeting (hCD19) or no CAR T cells. The graph shows the percentage of immune cells of all analyzed live cells. **I)** Sequencing plot showing the *Keap1* loss-of-function mutation R470H that was engineered in PDAC cells using base editing. The sequence of unedited PDAC cells (top) is shown in comparison to the sequence of a single cell clone with a homozygous R470H mutation (bottom). **G)** Experimental schematic: C57BL/6 mice received an orthotopic injection of 600,000 murine PDAC cells with or without *Keap1* loss-of-function mutation (R470H), followed by a subsequent intraperitoneal injection of freshly prepared mmeso CAR T cells (2 million) or control CAR T cells four days later (n=5 per experimental group). Mice were monitored until a humane endpoint was reached and their survival was recorded. **H)** A Kaplan-Meier curve showing survival of mice bearing *Keap1* R470H or control infected PDAC tumors treated with control or mmeso CAR-T cells. *Keap1* mutation significantly extends survival in combination with mmeso CAR T cell treatment (Mantel-Cox test p=0.0277).

Next, we investigated whether *Keap1* loss in tumor cells causes any changes in the tumor microenvironment that may contribute to their increased susceptibility to CAR T cell therapy. We injected hCD19+ PDAC cells with control (*Olfr402*) KO or *Keap1* KO cells orthotopically into C57BL/6 mice and treated them with hCD19 CAR T cells or no CAR T cells (**Figure 5D**). When we isolated and analyzed the tumors 12 days later, untreated *Keap1* KO tumors were significantly larger than untreated control KO tumors (**Figure 5E**) and contained significantly fewer CD45+ immune cells (**Figure 5F**), but an increased percentage of E2-Crimson+ PDAC cells (**Figure S13A**). This is in line with multiple studies highlighting *Keap1* KO tumors as “immune-cold”^61,71,72^. Analysis of hCD19 surface antigen expression levels revealed no differences between untreated control KO and *Keap1* KO tumors (**Figure S13B**). The proportions of immune cells (with the exception of Ly6C+ Ly6G-MDSCs) were largely similar between untreated control KO and *Keap1* KO tumors (**Figure S13C-L**), rather the overall number of immune cells, including CD3+ T cells, was reduced in *Keap1* KO tumors (**Figure S13C-D**). Conversely, hCD19 CAR T cell treatment increased the percentage of immune cells in both *Keap1* KO and control KO tumors compared to untreated tumors. A closer look at different immune subsets revealed no differences in T cell subsets, B cells, NK cells and myeloid cell populations, between hCD19 CAR T cell treated control KO and *Keap1* KO tumors, indicating similar levels of immune infiltration in response to CAR T cell therapy. Taken together, untreated *Keap1* KO PDAC tumors are immune-cold compared to control KO tumors, but nevertheless remain susceptible to CAR T cell therapy.

To define the tumor cell-intrinsic changes caused by *Keap1* KO in PDAC cells that may underlie their sensitivity to CAR T cell therapy, we performed bulk RNA transcriptome analysis on PDAC cells isolated from untreated *Keap1* KO and control KO tumors. Following alignment of sequencing reads (**Figure S14A**), we performed differential gene expression analysis on control KO tumor cells compared to *Keap1* KO tumor cells (**Figure S14B**). GSEA on these differentially expressed genes revealed a strong depletion in inflammatory response signaling, such as IFN-γ, IFN-α, and TNF-α signaling, in *Keap1* KO PDAC cells (**Figure S14C)**, similar to that previously observed with *Slc33a1* loss (**Figure S12C**). Furthermore, *Keap1* loss led to an enrichment in reactive oxygen species pathways signaling, consistent with increased oxidative stress caused by Nrf2-hyperactivation.

Given that both *Slc33a1* KO and *Keap1* KO can sensitize PDAC cells to CAR T cell therapy, we compared their bulk transcriptome data (**Figure S12 and S14A-C**). GSEA confirmed enrichment of the UPR in *Slc33a1* KO cells and enrichment in reactive oxygen species pathways signaling in *Keap1* KO cells, respectively, as some of the top divergent pathways between the two gene KO (**Figure S14E**). Thus, *Keap1* loss and *Slc33a1* loss appear to impose different intrinsic stresses in PDAC cells that converge on increased sensitivity to CAR T cell therapy.

### A patient-derived mutation in *KEAP1* sensitizes to CAR T cell therapy

Loss-of-function mutations in *KEAP1* are observed in *KRAS*-driven malignancies, including in LUAD and PDAC^73^, prompting us to hypothesize that patient-derived mutations could also sensitize tumors to CAR T cell therapy. To test this hypothesis, we used base editing to engineer the murine orthologue of the human tumor-derived *Keap1* R470H mutation^74^ in WT PDAC cells (**Figure 5G**). Importantly, mutations at *KEAP1* R470 are observed in PDAC patients and have been previously shown to enhance NRF2 stability and target gene activation^75–77^. Next, we orthotopically transplanted WT or *Keap1^R470H/R470H^* PDAC cells followed by treatment with mmeso or control CAR T cells (**Figure 5H**). Prior to their injection *in vivo*, we confirmed that mmeso CAR T cells were efficient at killing WT PDAC cells *in vitro* (**Figure S15A**) while producing high levels of IFN-γ (**Figure S15B**). Consistent with our hypothesis, we observed significant survival extension in mice harboring *Keap1^R470H/R470H^* PDAC tumors compared to non-mutant PDAC tumors treated with mmeso CAR T cells (p=0.0277; **Figure 5I**). These results show that human tumor-derived *Keap1* mutations can sensitize tumors to CAR T cell therapy.

## Discussion

CAR T cell therapies have shown limited success in PDAC clinical trials for reasons that remain poorly understood. Importantly, CAR T cell failure alone is not sufficient to explain the resistance of PDAC to this therapy, suggesting that tumor-intrinsic factors may also be important. Efforts to discover tumor-intrinsic mechanisms of resistance have been hampered by the lack of faithful pre-clinical model systems that are amenable to high-throughput genetic screens. Especially challenging are large scale CRISPR-Cas9 KO screens in PDAC models *in vivo* due to limited orthotopic engraftment rates which do not achieve appropriate sgRNA representation required for screening genome-scale CRISPR libraries. As such, *in vivo* screens in PDAC models published to date often used small libraries, have rarely been performed in orthotopic allografts, and mainly focused on the discovery of genes essential for PDAC growth^78–80^. Multiple *in vivo* screens have sought to identify genes mediating general immune evasion in PDAC and have highlighted a role for antigen presentation, IFN-γ pathway signaling and lipid metabolism^25,81^. Screening for CAR T cell response modulators in other tumor models has revealed roles for antigen loss^19–21^, impaired death receptor signaling in tumor cells^22^, and alterations in the IFN-γ pathway^23–26^. However, to date, no *in vivo* screens for mediators of CAR T cell response have been performed in solid tumors, including PDAC models.

Here, we established a fully immunocompetent, orthotopic PDAC model that can be used for multiplexed *in vivo* CRISPR screens in tumor cells to discover and validate mediators of resistance to CAR T cell therapy. We used a modular CRISPR library that is divided into individual, non-redundant pools^23^, enabling us to perform iterative screening of more than one third of the protein-coding genome and all KEGG pathway associated genes. Through this large-scale screening approach, metabolic genes, particularly *Slc33a1*, emerged as critical modulators of CAR T cell therapy response in PDAC cells.

Using scRNA transcriptome analysis as an orthogonal approach, we discovered that PDAC cells resistant to CAR T cell therapy display a distinct metabolic cell state, characterized by reduced *Nrf2* pathway activation, glutathione metabolism, oxidative phosphorylation, and ROS signaling. Consistent with this observation, we demonstrated that PDAC cells with high *Nrf2* pathway activity, a marker of oxidative stress, are hypersensitive to, and eliminated by, CAR T cell therapy *in vivo* (**Figure 6**). Under homeostatic conditions, the antioxidant transcription factor Nrf2 is kept inactive by the oxidative stress sensor Keap1 and is only released upon stress conditions^82–84^. Thus, cells with loss of *Keap1* function are known to experience Nrf2 hyperactivation, resulting in a redox imbalance that can promote tissue injury and cancer progression^85,86^. Our experiments revealed that activation of the *Keap1/Nrf2* pathway, either via endogenous activation or genetic ablation of *Keap1*, was sufficient to sensitize PDAC cells to CAR T cell therapy (**Figure 6**). Additionally, we show that a patient-relevant mutation in *Keap1* can similarly sensitize PDAC tumors to CAR T cell treatment *in vivo*, representing to our knowledge, the first cancer-patient derived tumor mutation shown to enhance tumor susceptibility to CAR T cell therapy.

**Figure 6:**
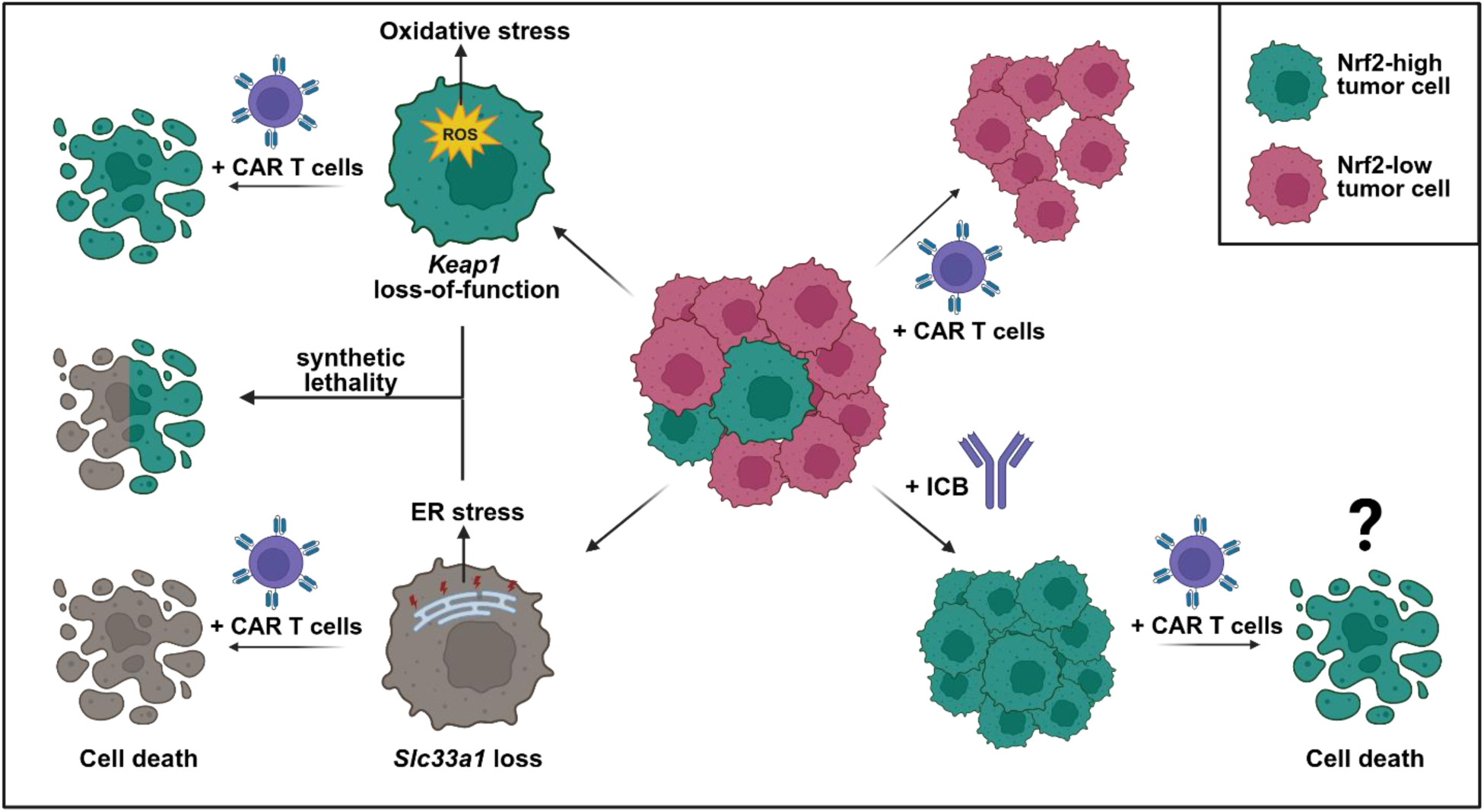
Model: Tumor cell-intrinsic stress states drive sensitivity to CAR T cell therapy. PDAC tumors contain *Nrf2*-high and *Nrf2*-low tumor cells. CAR T cell therapy effectively eliminates pre-existing *Nrf2*-high tumor cell populations. Additionally, tumor cell-intrinsic stress states, such as ER stress caused by *Slc33a1* loss, or oxidative stress caused by *Keap1* loss-of-function, sensitize to CAR T cell therapy. Combined inactivation of both *Slc33a1* and *Keap1* genes leads to tumor cell death driven by increases in oxidative stress and proteotoxic stress. Furthermore, the clinical observation that *Keap1* KO cells are particularly resistant to ICB therapy opens the possibility of the selection of *Nrf2*-hyperactivated tumor cells by ICB, which may then in turn be sensitized to CAR T cell therapy.

Intriguingly, our findings contrast with observations from ICB studies, where *Keap1* loss-of-function mutations have been associated with therapy resistance in pre-clinical models and LUAD patients^72,87,88^. *Keap1*-mutant tumors are characterized by an immunosuppressive, cold tumor environment, a finding that we recapitulated in our experiments^61,71,72^. Although both ICB and CAR T cell therapy fall under the immunotherapy umbrella, the two therapeutic modalities engage fundamentally different mechanisms. ICB relies on antibodies to enhance the endogenous immune response to antigens presented by the major histocompatibility complex I (MHC-I) on tumor cells, a process requiring both DCs and T cells^89^. In contrast, CAR T cell therapy does not require antigen presentation by MHC-I, nor the help of additional endogenous immune cell populations to mount an immune response. As such, our results may be reflective of mechanisms that are unique to adoptive cell-based immunotherapy and highlight the power of our modular *in vivo* screening platform to uncover such context-dependent tumor sensitivities. Furthermore, given that *Keap1* KO cells are particularly resistant to ICB therapy, this raises the intriguing possibility that ICB may select for a *Nrf2*-hyperactivated state that is responsive to subsequent CAR T cell therapy (**Figure 6**).

The acetyl-CoA transporter *Slc33a1*, identified as a determinant of resistance in our CRISPR-Cas9 screens, has recently been shown as a key mediator of tolerance to *Nrf2*-hyperactivation states in *Keap1* mutant LUAD^62^. Our data reinforce the important functional link between *Slc33a1* and the *Keap1*/*Nrf2* pathway. Indeed, combined loss of *Slc33a1* and *Keap1* in our PDAC cells resulted in synthetic lethality (**Figure 6**). We further showed that loss of *Slc33a1* alone sensitizes PDAC cells to CAR T cell therapy (**Figure 6**). Notably, *Slc33a1* loss in PDAC cells did not alter the tumor microenvironment, but caused tumor cell-intrinsic ER stress and the activation of the UPR, in agreement with prior studies in LUAD and pancreatic acinar cells^62,67^. Importantly, we show that this proteotoxic stress state is distinct from the oxidative stress state caused by *Keap1* loss, suggesting that disruption of distinct stress-adaptation pathways can independently mediate PDAC susceptibility to CAR T cell therapy.

Recently, the concept that tumor cell states can influence therapy response in PDAC has gained significant traction. For example, PDAC cells in the classical epithelial cell state have been shown to drive resistance to *KRAS* inhibitors, exhibiting enhanced sensitivity to cytotoxic T cell killing^33,34^. However, to date no tumor cell states have been discovered that influence CAR T cell therapy response. Our study decisively shows for the first time that tumor-intrinsic metabolic stress states can profoundly sensitize cancer cells to CAR T cell therapy. These cell states can be defined by oxidative stress, characterized by high activity of the *Nrf2* antioxidant response pathway, or proteotoxic stress, e.g. through *Slc33a1* loss.

Our data suggest that CAR T cell-sensitive cell states can arise through genetic and non-genetic mechanisms, including *Keap1* mutations and endogenous tumor evolution, respectively. Currently, no tumor-intrinsic mutations, except those present in tumor antigens, are known to modulate the response to CAR T cell therapy, and we lack any tumor biomarkers that predict CAR T cell response. Here, we show that perturbations driving PDAC development, evolution, and therapeutic resistance may also lead to a therapeutically exploitable collateral sensitivity to CAR T cell therapy. Future clinical studies are needed to identify if PDAC patients harboring tumor-intrinsic alterations in the *Keap1/Nrf2* antioxidant pathway are sensitive to CAR T cell therapy. Nevertheless, our findings open the possibility of therapeutic synergy through drugs targeting the *Nrf2* antioxidant pathway in combination with CAR T cell therapies to increase their efficacy in PDAC. Alternatively, therapies that drive the phenotypic selection for *Keap1*-mutant tumor cell states could be combined in sequence with CAR T cells that target this nascent vulnerability.

## Methods

### Cell culture and cell lines

The cell lines utilized in this study comprised murine PDAC cells (kindly provided by the Vander Heiden Laboratory), human embryonic kidney 293T cells (HEK293T; RRID:CVCL_0063) and Platinum-E (Plat-E; RRID:CVCL_B488) retroviral packaging cells. HEK293T cells, acquired from the American Type Culture Collection (ATCC), were sustained in DMEM with L-glutamine and sodium pyruvate (Corning, 10-013-CM), supplemented with 10% FBS. Murine PDAC cells were grown in DMEM with L-glutamine and sodium pyruvate (Corning, 10-013-CM), supplemented with 10% FBS. Plat-E cells were acquired from Cell Biolabs, Inc. and cultured in DMEM with 10% fetal bovine serum (FBS), 1 µg/mL puromycin, 10 µg/mL blasticidin, penicillin and streptomycin. Routine mycoplasma contamination checks were conducted for all cell lines.

Primary murine T cells were procured from mouse spleens and cultured on plates coated with activating antibodies, as described in the CAR-T cell production methods. The T cell medium consisted of RPMI with L-glutamine (Corning, 10-040-CM), supplemented with 10% FBS, recombinant human IL-2 (rhIL-2, final concentration of 20 ng/mL; Peprotech, Cat# 200-02-1mg), and 2-mercaptoethanol to a final concentration of 0.05 mM (Gibco, 21985023).

### Viral supernatant production

Lentiviral supernatant was produced using standard methods. Briefly, HEK293T cells were transfected with lentiviral transfer plasmid and packaging vector (psPAX2 and pMD2.G) using Mirus TransIT-LT1 (Mirus, MIR2305) as indicated by the manufacturer. For retroviral supernatant production, Plat-E cells were switched to DMEM + 10% FBS without antibiotics and transfected with retroviral transfer plasmid using Mirus TransIT-LT1 (Mirus, MIR2305) as indicated by the manufacturer. Viral supernatant was collected 48 hours and 72 hours after transfection, passed through a 0.45 μm filter, and stored at 4 °C for a maximum of one day.

### CAR-T cell production

Murine CAR T cells were produced as previously described^23^. Briefly, CD8^+^ T cells were isolated from the spleens of 14-week-old male or female C57BL/6 mice (Jackson Laboratory) using Miltenyi Biotec CD8a (Ly-2) MicroBeads for mouse (positive selection kit; Miltenyi, 130-117-044) and LS columns (Miltenyi, Cat# 130-042-401) as per the manufacturer’s instructions. The isolated T cells were cultured at 1×10^6^ cells/mL on 6-well plates coated with anti-murine CD3e and anti-murine CD28 activating antibodies (Bio X-Cell, BE0001-1 and BE0015-1) in T cell media.

After 24 hours, activated T cells were collected, counted, and resuspended at 0.5×10^6^ in a 1:1 mixture of fresh T cell media with viral supernatant supplemented with protamine sulfate to a final concentration of 10 μg/mL (MS Biomedicals, ICN19472910). The cells were spin-infected at 1000xg for 1.5 hours at 37 °C on new antibody-coated plates. The next day, T cells were again collected, counted, and resuspended at 1×10^6^ cells/mL in fresh T cell media, re-plated on new antibody-coated plates. Lastly, 24 hours later, T cells were collected, counted, and the percentage of CAR^+^ (GFP^+^) T cells determined by flow cytometry. The desired number of CAR T cells was then prepared for injection by resuspension in saline and injected via tail vein. Alternatively, T cells were resuspended at this step in T cell media and plated for *in vitro* cytotoxicity assays.

### *In vitro* cytotoxicity assay

Briefly, target cells were counted and co-cultured with or without CAR T cells at specified Effector-to-Target (E:T) ratios, accounting for the CAR-T transduction efficiency, in T cell media. After approximately 24 hours, the cell suspension was subjected to flow cytometry analysis to evaluate live/dead status (via DAPI stain), %hCD19^+^ cells, and %CD8^+^ cells. Flow cytometry was also employed to determine the densities of each cell type (CAR-T, target cell, non-transduced T cell). The resulting target cell densities in CAR-T-containing wells were normalized to those in control wells, seeded with the same number of target cells but with control CAR-T or non-transduced T cells. All flow cytometry experiments were conducted with a minimum of 10,000 live cells (via DAPI exclusion) and subsequent data analysis.

### Interferon gamma release ELISA assay

The enzyme-linked immunosorbent assay (ELISA) followed standard procedures. Briefly, supernatants from *in vitro* CAR-T cytotoxicity assays were collected and centrifuged to eliminate any contaminating cells. The quantification of IFNγ released by CAR T cells in the supernatant was performed using the DuoSet ELISA kit for mouse IFNγ (R&D systems, DY485). Nunc MaxiSorp flat-bottom plates (Thermo Fisher Scientific, 44-2404-21) were employed for the assay, conducted on a Tecan Infinite 200 Pro machine according to the manufacturer’s instructions. To maintain the assay within the linear range of the kit, the supernatant was initially diluted at 1:10 in reagent diluent. Subsequently, a minimum of six serial 4-fold dilutions were executed. For each plate, at least one standard curve was generated, and the entire experiment included at least two standard curves, constructed using standard solutions supplied by the manufacturer. The substrate solution used was 1-StepTM Ultra TMB-ELISA (Thermo Fisher Scientific, 34028), and the stop solution employed was 2N sulfuric acid (VWR, BDH7500-1). Bovine serum albumin (BSA; Sigma, A8022-500G) was prepared as a sterile-filtered 5% stock in PBS (Corning, 21-031-CV).

### Fluorescence-activated cell sorting and analysis

The BD LSRFortessa Cell Analyzer (RRID:SCR_018655) in tube or plate reader format, with BD FACSDiva Software (RRID:SCR_001456) v9.0 was used for data collection. All flow cytometry experiments were conducted with a minimum of 10,000 live cells (via DAPI exclusion). Downstream analysis was performed using FlowJo 10.9.0. The BD FACSAria II Cell Sorter (RRID:SCR_018934) was used for cell sorting.

### Western blotting

Cells were lysed with RIPA buffer (Boston BioProducts, BP-115) supplemented with 1X protease inhibitor mix (cOmplete EDTA-free, 11873580001, Roche). Protein concentration of cell lysates was determined using Pierce BCA Protein Assay (ThermoFisher Scientific, 23225). Total protein (25-40 μg) was separated on 4-12% Bis-Tris gradient SDS-PAGE gels (Life Technologies) and then transferred to PVDF membranes (IPVH00010, EMD Millipore) for blotting.

### Base editing

An sgRNA targeting R470 in *Keap1* was cloned into the pUSEBB backbone^90^. Lentivirus was produced and WT PDAC cells were transduced and sorted for sgRNA expression (BFP+). Next, cells were transduced with a cytosine base editing (CBE) reporter (mScarlet+)^90^, sorted and electroporated with *in vitro* transcribed mRNA encoding CBE6-NG. Successful base editing at the CBE reporter leads to GFP expression. After 72 hours of recovery, cells were sorted for GFP to enrich for base-edited PDAC cells. Next, these cells were single cell-cloned and successful editing was confirmed via Sanger sequencing. A single cell clone with homozygous *Keap1* R470H mutation was used for further experiments.

### Ethics statement

The research conducted in this study was conducted ethically and complies with all relevant guidelines and regulations. Animal studies were performed under strict compliance with the Committee of Animal Care (CAC) at the Massachusetts Institute of Technology (protocol number 2402000631).

### Animal housing and maintenance

All mouse experiments were conducted under Institutional Animal Care and Use Committee (IACUC)-approved animal protocols at the Massachusetts Institute of Technology (MIT). The mouse strains utilized in this study included C57BL/6 (The Jackson Laboratories; RRID:IMSR_JAX:000664) and NOD/SCID/IL-2Rg^−/−^ (NSG; The Jackson Laboratories; RRID:IMSR_JAX:005557), as indicated in the figure legends. All experimental mice used were 10-16 weeks old. Mice were housed under social conditions (two to five mice per cage) on a 12-hour dark/12-hour light cycle, ambient temperature 21 °C ± 1 °C, and humidity 50% ± 10%. All animals were housed in the pathogen-free animal facility of the MIT Koch Institute, in accordance with the animal care standards of the institutions. Food and water were provided *ad libitum*. All animal research at MIT is conducted under humane conditions with utmost regard for animal welfare. The animal care facility staff is headed by a chief veterinarian and includes a veterinary assistant, animal care technicians, and administrative support. MIT adheres to institutional standards for the humane use and care of animals, which have been established to assure compliance with all applicable federal and state regulations for the purchase, transportation, housing, and research use of animals.

### PDAC cell transplantation

PDAC cells were orthotopically injected into the pancreas as previously described^91^. Briefly, PDAC cells were resuspended in HBSS and 600,000 cells in 100 µl were orthotopically injected into the tail of the pancreas using a 26G1/2 needle. Successful injection was confirmed by visible bubble formation in the pancreatic parenchyma. Four days later, mice were intraperitoneally injected with CAR T cells suspended in saline on the opposing abdominal site.

For experiments in which tumor growth was monitored, mice were imaged every other day using ultrasound. As indicated, tumors were either isolated 10 days after initial injection or mice were closely monitored for signs of disease and distress, and sacrificed upon the appearance of morbidity, in accordance with CAC and DCM policy.

### Limiting GFP-dilution assay

600,000 PDAC cells were orthotopically transplanted into C57BL/6 mice. Five experimental groups of PDAC cells were tested containing diminishing amounts of GFP+ PDAC cells in an RFP+ PDAC cell population: (1) 50% RFP+ and 50% GFP+, (2) 99%RFP+ with 1% GFP+, (3) 99.9% RFP+ with 0.1% GFP+, (4) 99.95% RFP+ with 0.05% GFP+, (5) 99.975% RFP+ with 0.025% GFP+ PDAC cells.

Ten days after transplantation, PDAC tumors were isolated and analyzed by flow cytometry for GFP+ PDAC cells. The recovered percentage of GFP+ cells were then compared to the initial percentage of GFP+ PDAC cells prior to orthotopic transplantation.

### Competition assay

For *in vitro* competition assays, PDAC cells with or without hCD19 antigen were mixed at equal ratios and seeded at 1×10^4^ cells per well into a 48-well plate. Next, freshly prepared control (EGFRvIII) or hCD19 CAR T cells were added at an E:T ratio of 10:1 to the PDAC cells. After approximately 24 hours, the cell suspensions were subjected to flow cytometry analysis to evaluate live/dead status (via DAPI stain), %hCD19^+^ PDAC cells (GFP+) and %hCD19-PDAC cells (E2-Crimson+). he resulting target cell densities in CAR-T-containing wells were normalized to those in control wells, seeded with the same number of target cells but with control CAR-T or non-transduced T cells.

For *in vivo* competition assays, PDAC cells with or without hCD19 antigen were mixed at equal ratios and 600,000 total PDAC cells were orthotopically transplanted into C57BL/6 mice. Four days later, mice were injected with 2.5 million control (EGFRvIII) or hCD19 CAR T cells, or with saline as a negative control. Ten days after transplantation, PDAC tumors were isolated and analyzed by flow cytometry. Antigen-negative PDAC cells were identified by E2-Crimson expression and antigen-positive PDAC cells were identified by GFP expression. The ratios of GFP+ to E2-Crimson+ cells were compared between the three experimental groups.

### Pooled sgRNA screening

The previously described modular SKY CRISPR library which is divided into 48 individual, non-redundant pools that can be individually screened or combined into larger libraries was used^23^. In this library, Kyoto Encyclopedia of Genes and Genomes (KEGG (RRID:SCR_012773)) protein-coding genes are distributed into pools by pathway ranking, leading to the first 15 pools of the library covering all KEGG pathways. After the first 15 pools, the gene distribution is random. Thus, we elected to screen the first 16 pools of this library in pairs of two to cover all major signaling pathways. To preserve library complexity, a minimum of 1000-fold coverage of the sgRNA library was maintained at each *in vitro* step before the screen, and at a minimum of 150-fold coverage was maintained in all screens completed. Cloned and sequenced plasmid pools, and viral supernatant were generated by the Broad Institute’s GPP. For screens, Cas9+ cells were thawed, recovered, and expanded for 5 days to ensure robust growth, and then tested for cutting efficiency to ensure high rates of editing efficiency^92^. In brief, Cas9+ PDAC cells were transduced with the pXPR-047 plasmid that delivers both tdTomato and a sgRNA targeting tdTomato. Thus, active Cas9-expressing lines will result in a reduction in tdTomato over time. After cutting assays were completed, cells were expanded over three additional days and infected with sub-pools. For each of the 16 sub-pools, 6×10^6^ cells were spin-infected with predetermined amounts of viral supernatant such that 30-50% of all cells were infected (expressed E2-Crimson, and survived puromycin selection; MOI«1). Forty-eight hours later, 2 million transduced PDAC cells were started on puromycin selection (5 μg/ml, Gibco, A1113803). Cells were selected over seven days and allowed to recover for two days. On the day of injection into mice, the appropriate number of infected, selected, and recovered cells were combined so that each screen contained two pools. For each screen, PDAC cells injected into the pancreas of C57BL/6 mice as described under PDAC cell transplantation section of this manuscript. On the same day as the injection, input samples were collected. Four days later, 2.5 million control (EGFRvIII) or hCD19 CAR-T cells were adoptively transplanted into mice via intraperitoneal injection. Mice were sacrificed upon relapse and their tumors were isolated and dissociated into single cell suspensions. For this purpose, tumors were minced with a scalpel and incubated for 30 min in a HBSS-based digestion buffer containing 10 mM HEPES (Invitrogen, #15630080), 12.5U Collagenase IV (Millipore Sigma, C2674), and 200U DNaseI (NEB, #M0303) at 37°C. Next, tumors were further dissociated using a gentleMACS tissue dissociator (Miltenyi Biotec). Enzyme-activity was quenched with FBS and the samples were filtered through sequential filtering steps to obtain single cell solutions. Finally, gDNA from all cells was isolated using the Machery Nagel L Midi NucleoSpin Blood Kit (Clontech, 740954.20). Modifications to the manufacturer’s instructions were added as follows: in step1, cells were lysed in the kit’s proteinase K containing lysis buffer for longer (overnight at 70 °C). The next morning, lysates were allowed to cool to room temperature, 4.1 μl of RNase A (20 mg/mL; Clontech, 740505) was added, and cells were incubated for 5 minutes at room temperature. The procedure then continued as indicated by the manufacturer. PCR inhibitors were removed from the resulting gDNA (Zymo Research, D6030) and the concentration of the resulting gDNA was measured using the Qubit dsDNA HS assay kit (ThermoFisher, Q32854), and if necessary, diluted to 200 ng/μl with elution buffer. gDNA was then submitted for Illumina sequencing. Data were analyzed as specified in the Data Analysis section of this manuscript.

### Single cell transcriptome profiling

C57BL/6 mice were orthotopically injected with 6×10^5^ hCD19+ PDAC cells, followed by intraperitoneal injection with murine CAR-T cells targeting either hCD19 or a control epitope (human EGFRvIII). CAR T cell treated tumors were isolated after 10 days for control CAR T cell treated mice and after 15 days for hCD19 CAR T cell treated mice, at which stage all PDAC tumors had reached a similar weight. Tumors were dissociated into single cells and 10,000 live cells were submitted for transcriptional profiling. Single-cell expression libraries were prepared using the 10x Genomics Chromium v3 reagents. Data were analyzed as specified in the Data Analysis section of this manuscript.

### Bulk tumor cell transcriptome profiling

C57BL/6 mice were orthotopically injected with 6×10^5^ hCD19+ PDAC cells with a control (*Olfr402*) KO, *Slc33a1* KO or *Keap1* KO. Tumors were isolated after 12 days, dissociated into single cells and sorted for E2-Crimson+ tumor cells. At least 100,000 tumor cells per sample were submitted in DNA/RNA shield solution (Zymo Research, #R1100) to Plasmidsaurus for bulk RNA transcriptome profiling. Data were analyzed by Plasmidsaurus as specified in the Data Analysis section of this manuscript.

### *Nrf2* reporter experiment

WT PDAC cells were transduced with a previously described *Nrf2* 8xARE-GFP reporter for which GFP expression indicates functional *Nrf2* transcriptional activity^58^. For *in vitro* validation of reporter function, *Nrf2* reporter PDAC cells were treated with tert-butylhydroquinone (tBHQ) and analyzed by flow cytometry for GFP expression 24 hours later. For in vivo experiments, C57BL/6 mice were orthotopically injected with 6×10^5^ *Nrf2* reporter PDAC cells, followed by intraperitoneal injection with murine CAR-T cells targeting either murine mesothelin or a control epitope (human EGFRvIII). CAR T cell treated tumors were isolated after 10 days, dissociated and PDAC cells (BFP+) were analyzed for GFP expression by flow cytometry.

### Tumor flow cytometry profiling

C57BL/6 mice were orthotopically injected with 6×10^5^ hCD19+ PDAC cells with a control (*Olfr402*) KO, *Slc33a1* KO or *Keap1* KO, followed by intraperitoneal injection with murine CAR-T cells targeting either hCD19 or a control epitope (human EGFRvIII). CAR T cell treated tumors were isolated after 12 days, and dissociated into single cells (as described above) for flow cytometric analysis. Samples were blocked with Fcr blocking reagent for mouse (Miltenyi Biotec 130-092-575) for 10 minutes at 4°C. Then samples were resuspended in antibody solution and incubated at 4°C for 1 hour. Following a wash step with PBS, samples were fixed using fresh 4% PFA for 20 minutes at room temperature. After another wash, samples were resuspended in PBS and data were collected within 24 hours using the BD LSRFortessa Cell Analyzer (RRID:SCR_018655) in tube or plate reader format, with BD FACSDiva Software (RRID:SCR_001456) v9.0. Downstream analysis was performed using FlowJo 10.9.0.

### Tumor histology

PDAC tumors were fixed with 10% Formalin for 24 hours at 4°C and subsequently embedded in paraffin. Tumors were sectioned into 4 µm thick sections and stained with hematoxylin and eosin (H&E). Tumor sections were imaged using an Aperio Digital Pathology Slide scanner (Leica Biosystems) at 20x resolution and analyzed using the corresponding Aperio ImageScope software (RRID:SCR_020993).

### Statistical analysis

All statistical analyses were conducted using GraphPad Prism 10 (GraphPad Software Inc (RRID:SCR_002798)). The specific statistical tests performed for each analysis are outlined in the figure legends. Differences were considered statistically significant for P-values ≤ 0.05.

## Antibodies

### Western blotting

anti-β-ACTIN (Cell Signaling Technology Cat# 4967, RRID:AB_330288), anti-Cas9 (Active Motif Cat# 61577, RRID:AB_2793684), anti-Slc33a1 (Sigma-Aldrich Cat# WH0009197M7, RRID:AB_2286124), anti-Nrf2 (Cell Signaling Technology Cat# 12721, RRID:AB_2715528), anti-Keap1 (Cell Signaling Technology Cat# 8047, RRID:AB_10860776), anti-rabbit IgG HRP-linked antibody (Cell Signaling Technology Cat# 7074, RRID:AB_2099233), anti-mouse IgG HRP-linked antibody (Cell Signaling Technology Cat# 7076, RRID:AB_330924), anti-rat IgG HRP-linked antibody (Cell Signaling Technology Cat# 7077, RRID:AB_10694715), Rabbit anti-Armenian Hamster IgG H&L (HRP) (Abcam Cat# ab5745, RRID:AB_955407).

### Flow cytometry

anti-human CD19-BV785 (BioLegend Cat# 302240, RRID:AB_2563442), anti-mouse Cd8a-PE/Cy7 (BioLegend Cat# 100722, RRID:AB_312761), anti-murine Cd45-PerCP-Cy5.5 (BioLegend Cat# 103132, RRID:AB_893340), anti-murine Cd45-BV785 (BioLegend Cat# 103149, RRID:AB_2564590), anti-murine Cd45-BV711(BioLegend Cat# 103147, RRID:AB_2564383), anti-murine Cd11b-APC-Cy7 (BioLegend Cat# 101226, RRID:AB_830642), anti-murine Cd11c-PerCP Cy5.5 (BioLegend Cat# 117328, RRID:AB_2129641), anti-murine Ly6C-BV605 (BioLegend Cat# 128035, RRID:AB_2562352), anti-murine Ly6G-FITC (Thermo Fisher Scientific Cat# 11-9668-82, RRID:AB_2572532), anti-murine F4/80-SparkUV387 (BioLegend Cat# 285504, RRID:AB_3717064), anti-murine Cd19-785 (BioLegend Cat# 115543, RRID:AB_11218994), anti-murine Nk1.1-PECy7 (Thermo Fisher Scientific Cat# 25-5941-81, RRID:AB_469664), anti-murine CD3-SparkUV387 (BioLegend Cat# 100284), anti-murine Cd8a-APC (BioLegend Cat# 100712, RRID:AB_312751), and anti-murine Cd4-PECy7 (BioLegend Cat# 100421, RRID:AB_312706).

## Plasmid constructs

Plasmid constructs:
| Plasmid | Producer |
| --- | --- |
| psPAX2 (lentiviral packaging) | RRID:Addgene_12260 |
| pMD2.G (VSV-g envelope) | RRID:Addgene_12259 |
| lentiCas9-Blast | RRID:Addgene_52962 |
| hCD19 CAR-P2A-GFP plasmid | Ramos, Koch and Liu et al., Nature Communications, 2023 |
| EGFRvIII CAR-P2A-GFP plasmid | Ramos, Koch and Liu et al. Nature Communications, 2023 |
| pMMLV hCD19(-ICD) | Fröse and Rowley et al., Science Advances, 2024 |
| mmeso CAR-P2A-GFP plasmid | Sequence generously shared by the June lab and then cloned into the same plasmid backbone as other CAR plasmids used in this study. |
| pRDACrimson_170 | Ramos, Koch and Liu et al., Nature Communications, 2023 |
| pXPR-047 | RRID:Addgene_107145 with tdTomato instead of eGFP; |
| *Nrf2* 8xARE-GFP reporter | Zucker et al., Cell, 2025 |
| pLEX307 | RRID:Addgene_41392; backbone used for expression of Cas9 resistant *Slc33a1* for overexpression |
| pUSEBB | Sánchez-Rivera and Diaz et al., Nature Biotechnology, 2022 |
| Cytosine base editor (CBE) reporter | RRID:Addgene_136895 |
| SpCas9 CBE6-IVT template | RRID:Addgene_215834 |

### CRISPR plasmids

For hit validation, the top 2 scoring sgRNAs for each gene were taken from the SKY library. All other sgRNAs were designed using the Broad Institute’s sgRNA Designer tool (https://portals.broadinstitute.org/gppx/crispick/public). sgRNAs were cloned into pRDACrimson_170 or pUSEBB to generate KOs of indicated genes. The sgRNA sequence for *Keap1* R470H was kindly provided by Acosta, Johnson and Gould et al.^74^ and is a murine equivalent of the human *KEAP1* R470H mutation.

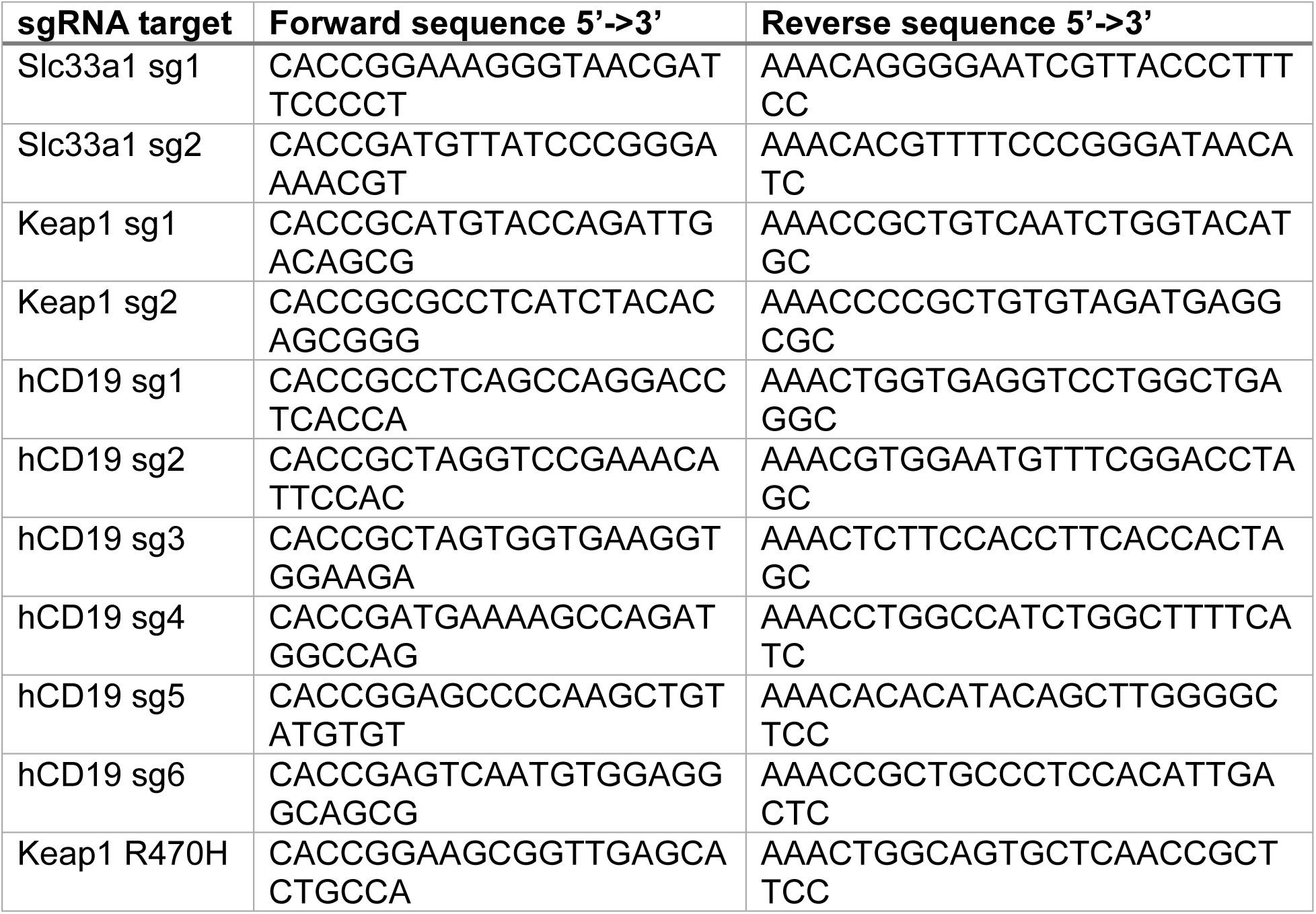

## Data Analysis

### Primary CRISPR-Cas9 KO Screen

Count data was assigned to SKY library barcodes by the Broad Institute’s Genomic Perturbation Platform (GPP) using PoolQ (version 3.3.1) from the output of 5 separate sequencing runs. The resulting 24 counts files were manually assembled using Tibco Spotfire version 11.4.1.16 and R version 4.4.1^93^ with tidyverse version 2.0.0^94^ and exported to 8 data matrices. Each matrix contains data for a pair of pools of the SKY library and contains sgRNA identifiers, the corresponding gene association for these identifiers and count data for input, EGFRvIII, and CD19 replicates. Interquartile ranges were calculated for each sample and 5 outliers were identified and excluded from analysis, these were: pool 1+2, EGFRvIII replicate 2, pool 3+4 EGFRvIII replicate 5, pool 7+8 CD19 replicate 16, and pool 13+14 replicates CD19 14 and EGFRvIII 3. These count matrices were then used as input for differential comparison with MAGeCK test version 0.5.9.4^39^ with the parameters adjust-method fdr, gene-lfc-method median and norm-method control using a list of negative control guides that target intergenic or olfactory gene sequences. After testing, guide and gene-level data results for each experiment and pool were imported, processed and visualized in R using MAGeCKFlute version 2.8.0^95^ and tidyverse 2.2.0^94^.

Gene and guide-level data were then aggregated into single data frames in order to visualize differential comparisons across the pools. The 16 pools of the SKY library screened in this study contains 29677 guides targeting barcodes. Of these 27705 target a single gene for a total of 7154 screened genes. Genes of interest and those exceeding different log fold change and FDR thresholds were then selected for validation screening.

### Validation CRISPR-Cas9 KO Screen

A validation library was prepared consisting of 1960 sgRNAs. These include 376 non-targeting control sgRNAs and sgRNAs that target 328 genes, usually with 6 distinct guides. Differential analysis was performed for validation screen data with MAGeck test in the same way as for the primary screen. All validation screen samples were used in analysis. Metabolic genes were defined as those encoding enzymes or transporters participating in canonical metabolic pathways based on KEGG, Reactome, and GO. Genes involved in nutrient sensing, autophagy, and metabolic stress adaptation were classified as metabolism-adjacent.

### Single cell transcriptome profiling

Sequencing data was aligned to the Mus_musculus.GRCm38v98.chr reference genome (with human CD19, Cas9 and EGFP added to it) and converted to fastq files using bcl2fastq (v2.20.0.422). Cell count matrices were generated using cellranger (v.6.1.1). Matrices were analyzed by Seurat (v4.0.4) for R (v4.0.2). Digital gene expression matrices were filtered on the following criteria to exclude low-quality cells: number of genes detected in each cell **(=**nFeature_RNA >100, <8000), the total number of molecules detected within a cell **(=**nCount_RNA; top 10% removed) and percent mitochondrial reads (=percent.mito; <25%). The data were further filtered to exclude ambient RNA (SoupX) and doublets (DoubletFinder). Next, data from all tumors was combined into a shared data object and was normalized, scaled, and principal component analysis was performed on the scaled data. The standard deviation of principal components was quantified using an elbow plot, and input dimensions for SNN clustering (20) at which standard deviation = 2 was used. SNN clustering was performed to generate UMAP plots (k.param = 20, res = 0.8). Unique markers for each cluster were identified using FindAllMarkers function. Clusters were annotated using a mix of manual annotation based on well-known lineage genes, such as *Cd68* for macrophages, and the Bioconductor package SingleR. Tumor cells were identified by hCD19 and Cas9 expression and subsetted. Following normalization and scaling, principal component analysis was performed on the scaled data. SNN clustering was performed to generate UMAP plots (k.param = 10, res = 0.4). Differential abundance analysis was performed using MiloR to compare UMAP changes between control and hCD19 CAR T cell treated PDAC cells^96^. Differentially expressed genes between PDAC cell with control CAR T cell treatment and PDAC cells with hCD19 CAR T cell treatment were identified using the FindMarkers function. The top 100 differentially expressed genes were used for Gene Ontology (RRID:SCR_002811) analysis. A ranked list based on all differentially expressed genes was used for Gene Set Enrichment Analysis (RRID:SCR_003199). For the generation of a murine Nrf2-responsive gene signature, the human NRF2 WikiPathway gene set (species = “human”, collection = “C2”, sub collection = “CP:WIKIPATHWAYS”, gene set= “M39454”) was used and murine orthologs to all genes were identified using the “orthologs” function of the R package “Disco” (v 0.6). The resulting gene list was filtered to only contain genes expressed in our PDAC cells. The murine Nrf2-responsive gene signature was then created of the resulting genes using the Seurat function “AddModuleScore” and displayed in a Feature Plot.

### Bulk tumor cell transcriptome profiling

Quality of the fastq files was assessed using FastQC v0.12.1. Reads were then quality filtered using fastp v0.24.0 with poly-X tail trimming, 3’ quality-based tail trimming, a minimum Phred quality score of 15, and a minimum length requirement of 50 bp. Quality-filtered reads were aligned to the reference genome using STAR aligner v2.7.11 with non-canonical splice junction removal and output of unmapped reads, followed by coordinate sorting using samtools v1.22.1. PCR and optical duplicates were removed using UMI-based deduplication with UMIcollapse v1.1.0. Alignment quality metrics, strand specificity, and read distribution across genomic features were assessed using RSeQC v5.0.4 and Qualimap v2.3, with results aggregated into a comprehensive quality control report using MultiQC v1.32. Gene-level expression quantification was performed using featureCounts (subread package v2.1.1) with strand-specific counting, multi-mapping read fractional assignment, exons and three prime UTR as the feature identifiers, and grouped by gene_id. Final gene counts were annotated with gene biotype and other metadata extracted from the reference GTF file. Sample-sample correlations for sample-sample heatmap and PCA were calculated on normalized counts (TMM, trimmed mean of M-values) using Pearson correlation. Differential expression was done with edgeR v4.0.16 using standard practice including filtering for low-expressed genes with edgeR::filterByExprwith default values. Functional enrichment was performed using gene set enrichment analysis with gseapy v0.12 using the MSigDB Hallmark gene set.

## List of Supplementary Materials

Fig. S1 to S15 for multiple supplementary figures, as well as supplementary tables 1 to 4.

## Data availability

All data are available with this manuscript. Single cell RNA sequencing data and bulk RNA sequencing data are available at Gene Expression Omnibus (GEO; RRID:SCR_005012) under GSE304184. All raw screen data are also accessible at the following GitHub (RRID:SCR_002630) repository: https://github.com/KochInstitute-Bioinformatics/Frose_SolidTumor_CART.

## Acknowledgments

We express gratitude to Koch Institute’s Robert A. Swanson (1969) Biotechnology Center, particularly the Massachusetts Institute of Technology Koch Institute Preclinical Modeling Core Facility (RRID:SCR_017899), the Massachusetts Institute of Technology Koch Institute Flow Cytometry Core Facility (RRID:SCR_017892), the Massachusetts Institute of Technology Koch Institute Histology Core Facility (RRID:SCR_019177), and the Massachusetts Institute of Technology Koch Institute Bioinformatics and Computing Core Facility (RRID:SCR_017894) for their technical assistance. CAW is partially supported by Cancer Center Support (core) Grant P30-CA14051 from the NCI to the Barbara K. Ostrom (1978) Bioinformatics and Computing Core Facility of the Swanson Biotechnology Center. We thank Professor Carl H. June for generously sharing the murine mesothelin CAR T cell sequence with us. We also thank Samuel I. Gould for kindly sharing the *Nrf2* 8xARE-GFP reporter with us. We are very grateful to Dr. Allison Lau and Dr. Sharanya Sivanand for sharing PDAC KPC cell lines with us. We would also like to thank Gabriel A. Camaaño Lasanta and Loona Meyer for technical assistance. Lastly, we would like to acknowledge Biorender (RRID:SCR_018361) for experimental layout figures in this manuscript.

## Author contributions (CRediT)

□ **Writing – original draft:** Julia Fröse

□ **Conceptualization:** Julia Fröse, Michael Hemann

□ **Investigation:** Julia Fröse, Evelyn Chen, Charles A. Whittaker, Paul Leclerc, Adam Langenbucher, Sean Doherty, Jasmine Shao, Riley D. Hellinger, Daniel Goulet

□ **Writing – review and editing:** Julia Fröse, Evelyn Chen, Charles A. Whittaker, Paul Leclerc, Adam Langenbucher, Riley D. Hellinger, Daniel Goulet, Jasmine Shao, Francisco Sánchez-Rivera, Tuomas Tammela, Michael Hemann

□ **Methodology:** Julia Fröse, Evelyn Chen, Michael Hemann

□ **Validation:** Julia Fröse, Evelyn Chen, Charles A. Whittaker, Paul Leclerc, Sean Doherty, Adam Langenbucher, Michael Hemann, Tuomas Tammela

□ **Formal analysis:** Julia Fröse, Charles A. Whittaker, Paul Leclerc, Evelyn Chen, Adam Langenbucher

□ **Project administration:** Julia Fröse, Michael Hemann

□ **Visualization:** Julia Fröse, Charles A. Whittaker

□ **Data curation:** Julia Fröse, Evelyn Chen, Charles A. Whittaker, Paul Leclerc, Adam Langenbucher

□ **Funding acquisition:** Michael Hemann, Francisco Sánchez-Rivera, and Tuomas Tammela

□ **Supervision:** Julia Fröse, Michael Hemann

