## Supplemental figures S1-S15 for "Tumor cell-intrinsic stress states drive sensitivity to CAR T cell-therapy in pancreatic cancer"

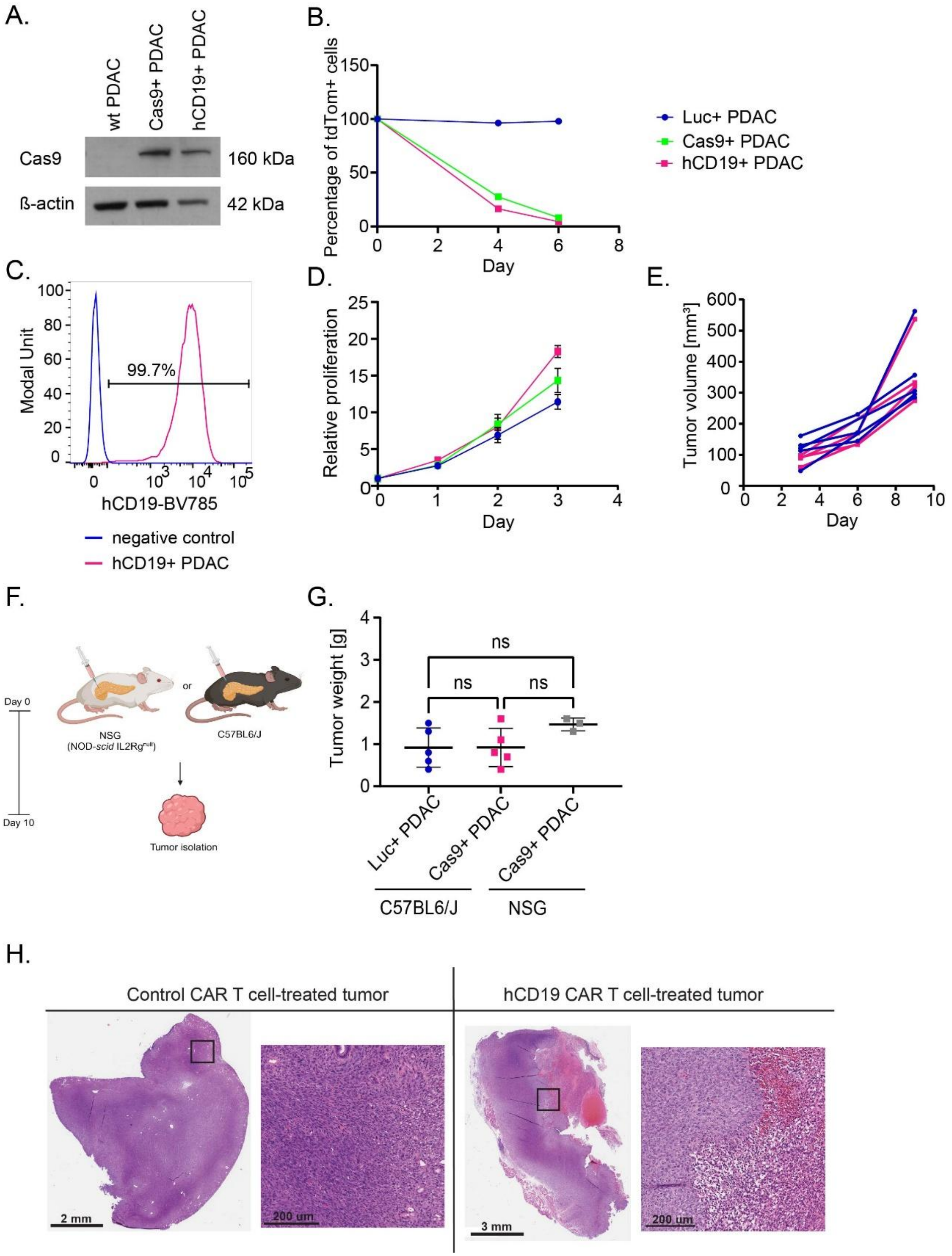


**Figure S1: Development of a CRISPR-Cas9 knock-out screening-amenable PDAC model for studying CAR T cell therapy resistance. A)** A western blot showing Cas9 protein expression in KPC PDAC cell lines. **B)** A graph showing Cas9 activity in PDAC cells *in vitro* as measured by disruption of tdTomato expression with an tdTomato-targeting sgRNA over a time period of 6 days. **C)** A flow cytometry plot showing hCD19 cell surface expression on hCD19+ PDAC cell compared to a negative control cell line. **D)** A graph showing *in vitro* proliferation of the Luc+ PDAC cell, Cas9+ PDAC cells, and hCD19+ PDAC cells. **E)** A graph showing *in vivo* proliferation of PDAC cells with and without Cas9 expression following orthotopic injection of 600,000 PDAC cells into C57BL/6 mice. **F)** Experimental schematic: Experimental schematic: C57BL/6 and NSG (Nod-scid IL2Rg$^{null}$) mice received an orthotopic injection of 600,000 murine PDAC cells with or without Cas9 and hCD19 expression. Tumors were isolated ten days later and their weight was measured. **G)** A graph showing the weight of PDAC tumors formed by Luc+ PDAC cells or hCD19+ PDAC cells in C57BL/6 mice and NSG mice. The tumor weights are not significantly different between mouse strains and PDAC cell lines. Statistical analysis with Ordinary One-Way ANOVA test (ns > 0.05). **H)** H&E-stained tumor sections from hCD19+ PDAC tumors treated with control CAR T cells targeting an irrelevant antigen (EGFRvIII) or treated with hCD19 CAR-T cells. PDAC tumors treated with hCD19 CAR-T cells show large necrotic areas and immune cell infiltration.

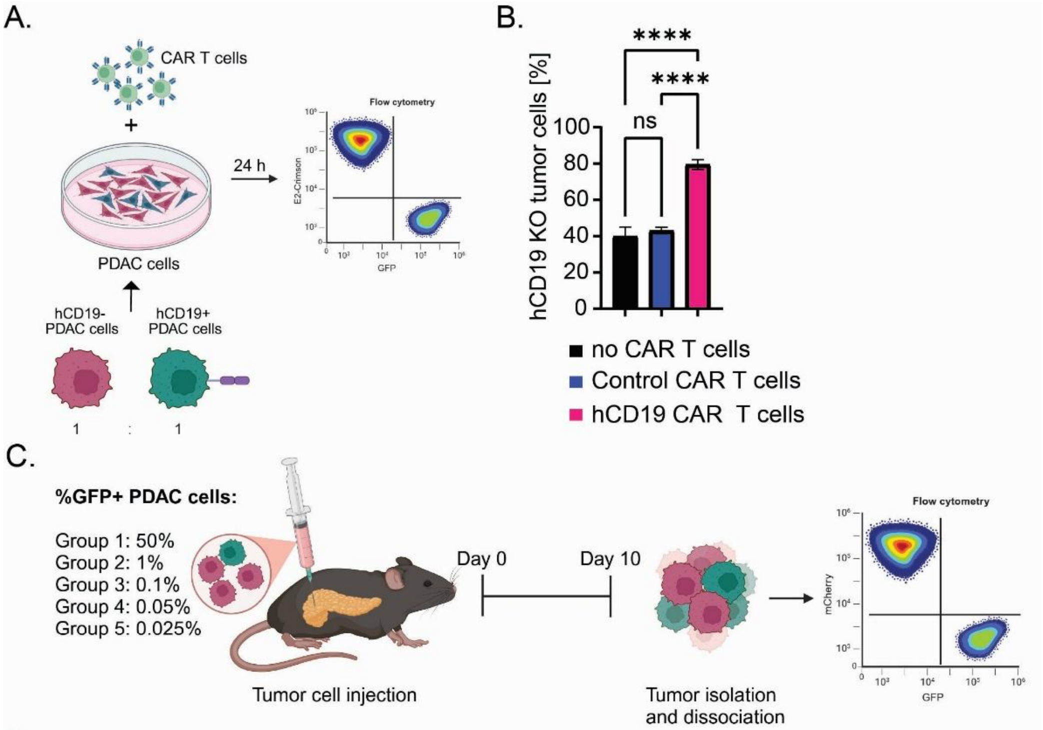


D.

| | | Expected input | | True Input | Output |
|---|---|---|---|---|---|
| Group | Total no. of PDAC cells | %GFP | Maximum sgRNA library size | %GFP | Average %GFP |
| 1 | 6.00E+05 | 50 | 2 | 51.5267% | 52.0774% |
| 2 | 6.00E+05 | 1 | 100 | 1.1974% | 1.1089% |
| 3 | 6.00E+05 | 0.1 | 1000 | 0.1058% | 0.0947% |
| 4 | 6.00E+05 | 0.05 | 2000 | 0.0453% | 0.1283% |
| 5 | 6.00E+05 | 0.025 | 4000 | 0.0268% | 0.0299% |

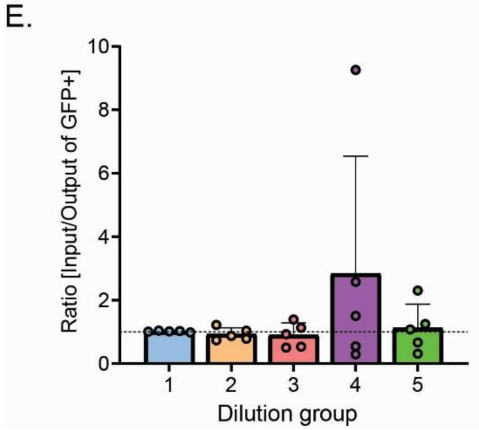


**Figure S2: A PDAC model amenable for *in vivo* CRISPR-Cas9 KO screening. A)** Experimental schematic: Testing the selective pressure conferred by CAR T cell treatment *in vitro* in a CAR T cell cytotoxicity assay. PDAC cells with hCD19 antigen expression (GFP+) were mixed at a 1:1 ratio with hCD19-negative PDAC cells (E2-Crimson+). Next, 10,000 PDAC cells from this mixture were incubated with 100,000 hCD19 CAR T cells or control (EGFRvIII) CAR T cells for 24 hours, followed by flow cytometric analysis. **B)** A bar graph showing the change in the percentage of hCD19-negative cells in a mixture of antigen-positive and -negative PDAC cells after CAR T cell treatment. Only treatment with hCD19 CAR T cells causes enrichment of antigen-negative PDAC cells. All conditions were analyzed in triplicate and error bars represent standard deviation. Statistical analysis with Ordinary One-Way Anova test (ns > 0.05, * ≤ 0.05, ** p ≤ 0.01, *** p ≤ 0.001, **** p ≤ 0.0001). **C)** Experimental schematic: Determining the maximal CRISPR library size that can be represented in a murine PDAC tumor model *in vivo* using a GFP dilution assay. Diminishing amounts of GFP+ hCD19+ PDAC cells were spiked into an mCherry+ hCD19+ PDAC cell population at the indicated percentages. A total of 600,000 hCD19+ PDAC cells was then orthotopically transplanted into C57BL/6 mice. In total, five experimental groups containing hCD19+ PDAC cells with GFP percentages ranging from 50% to 0.025% were transplanted into mice (n=5 per experimental group). After ten days, tumors were isolated, dissociated into single cell solutions and analyzed for mCherry+ and GFP+ tumor cells by flow cytometry. The percentage of recovered GFP+ cells (output) was then analyzed and compared to the initial GFP+ tumor cell percentage prior to injection (input). **D)** Table showing the five experimental groups and their inferred representation of CRISPR library size. The largest CRISPR library that can be represented in this hCD19+ PDAC tumor model is 4000 sgRNAs (150x representation) in a total of 600,000 tumor cells. **E)** A bar graph showing the ratio of initial percentage of GFP+ PDAC cells (input) to the GFP+ PDAC cell percentage at the experimental endpoint (output). A ratio ≥1 indicates that the GFP percentage remained stable throughout the entirety of the experiment.

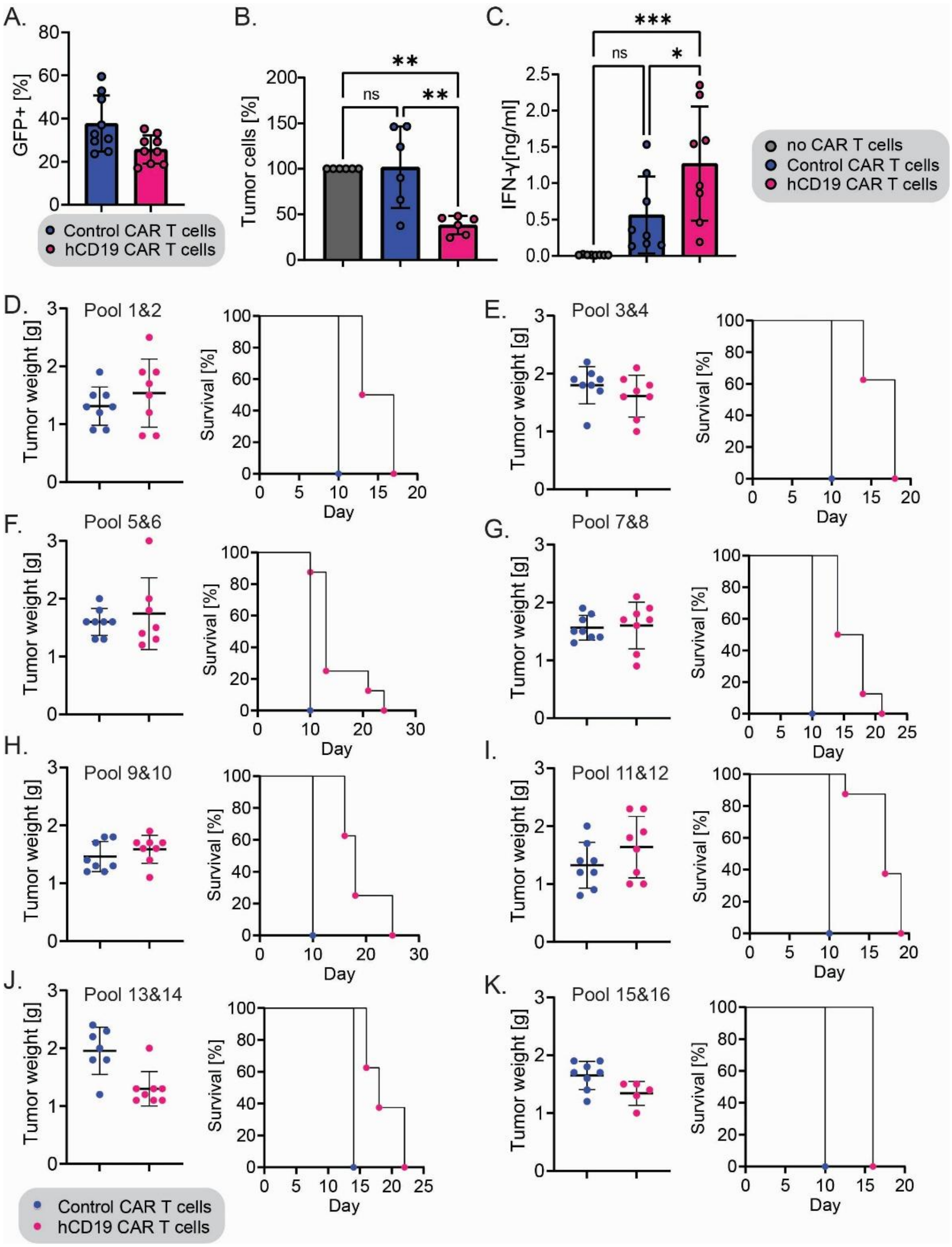


**Figure S3: CAR T cell quality control data and tumor data from eight independent CRISPR-Cas9 KO screens. A)** A bar graph showing the average transduction rate of murine CD8+ T cells with CAR-GFP constructs. **B)** A bar graph showing the average CAR T cell killing across six independent CAR T cell cytotoxicity assays involving incubation of hCD19 CAR T cells or control (EGFRvIII) CAR T cells with PDAC cells (E:T ratio = 10:1) for 24 hours. All conditions were analyzed in triplicate. Error bars represent standard deviation. Statistical analysis with Ordinary One-Way ANOVA test (ns > 0.05, * ≤ 0.05, ** p ≤ 0.01, *** p ≤ 0.001, **** p ≤ 0.0001). **C)** A bar graph showing the average IFN-γ release by hCD19 or control CAR T cells during the CAR T cell cytotoxicity assays across six independent experiments. All conditions were analyzed in triplicate and error bars represent standard deviation. Statistical analysis conducted using Ordinary One-Way ANOVA test (ns > 0.05, * ≤ 0.05, ** p ≤ 0.01, *** p ≤ 0.001, **** p ≤ 0.0001). **D-K)** Average tumor weights (left) and tumor day harvests (right) shown in Kaplan-Meier format for each independent CRISPR-Cas9 KO screen. D) Screen 1 with CRISPR library pools 1&2, E) Screen 2 with CRISPR library pools 3&4 , F) Screen 3 with CRISPR library pools 5&6, G) Screen 4 with CRISPR library pools 7&8, H) Screen 5 with CRISPR library pools 9&10, I) Screen 6 with CRISPR library pools 11&12, J) Screen 7 with CRISPR library pools 13&14, K) Screen 8 with CRISPR library pools 15&16.

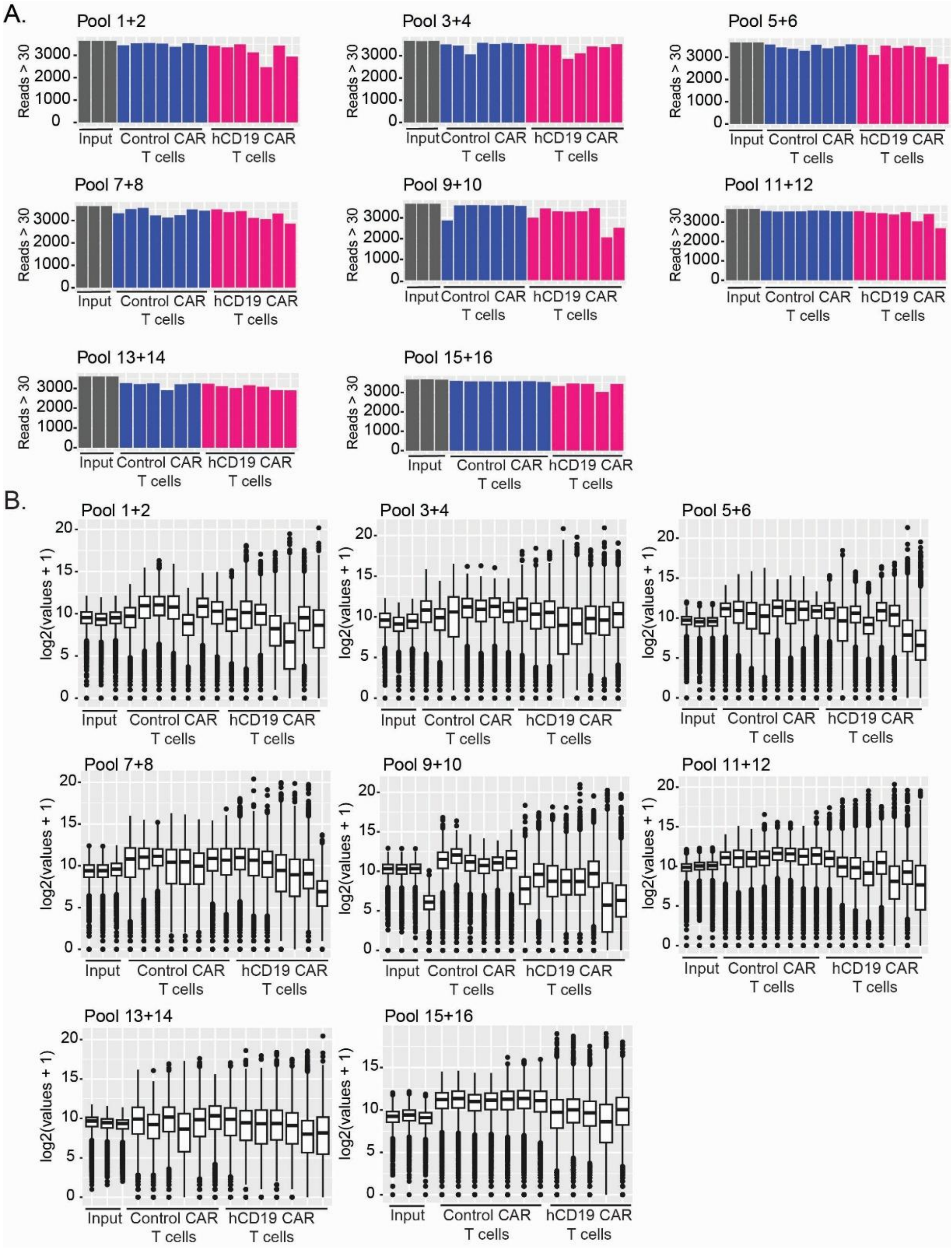


**Figure S4: Analysis of eight independent *in vivo* CRISPR-Cas9 KO screens. A)** A bar graph showing the number of represented sgRNA per mouse for each independent CRISPR-Cas9 KO screen. The number of sgRNAs with at least 30 reads is shown for PDAC cells containing the CRISPR libraries on the day of injection (input) compared to tumor cells isolated from mice treated with hCD19 CAR T cells or treated with control CAR T cells. **B)** Box plots showing the distribution of raw sgRNA counts in PDAC cells containing the CRISPR libraries on the day of injection (input) compared to tumor cells isolated from mice treated with hCD19 CAR T cells or treated with control CAR T cells for each independent CRISPR-Cas9 KO screen.

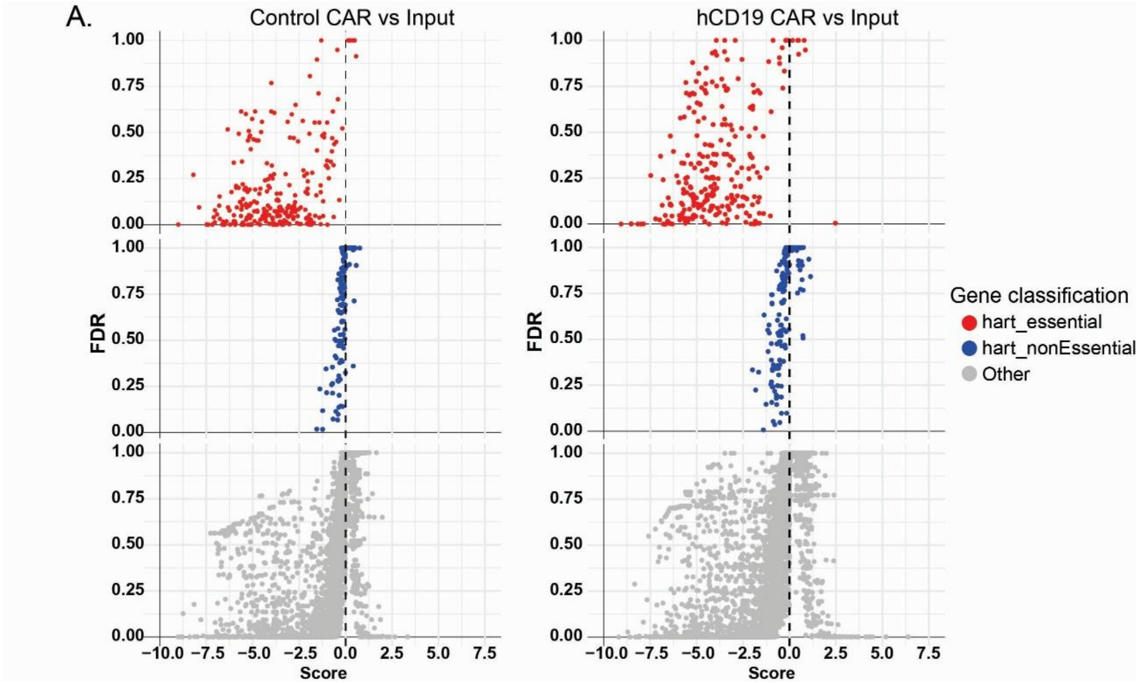


**Figure S5: Essential genes show predicted behavior *in vivo*.** Scatter plot highlighting the behavior of sgRNAs targeting genes classified as essential or non-essential, as defined by Hart et. al, compared to sgRNAs targeting all remaining genes in the first CRISPR-Cas9 KO screen (Pool 1 & 2) in control CAR T cell treated and hCD19 CAR T cell treated mice compared to tumor cells on the day of injection (input). FDR = False discovery rate, Score = Log2 fold change.

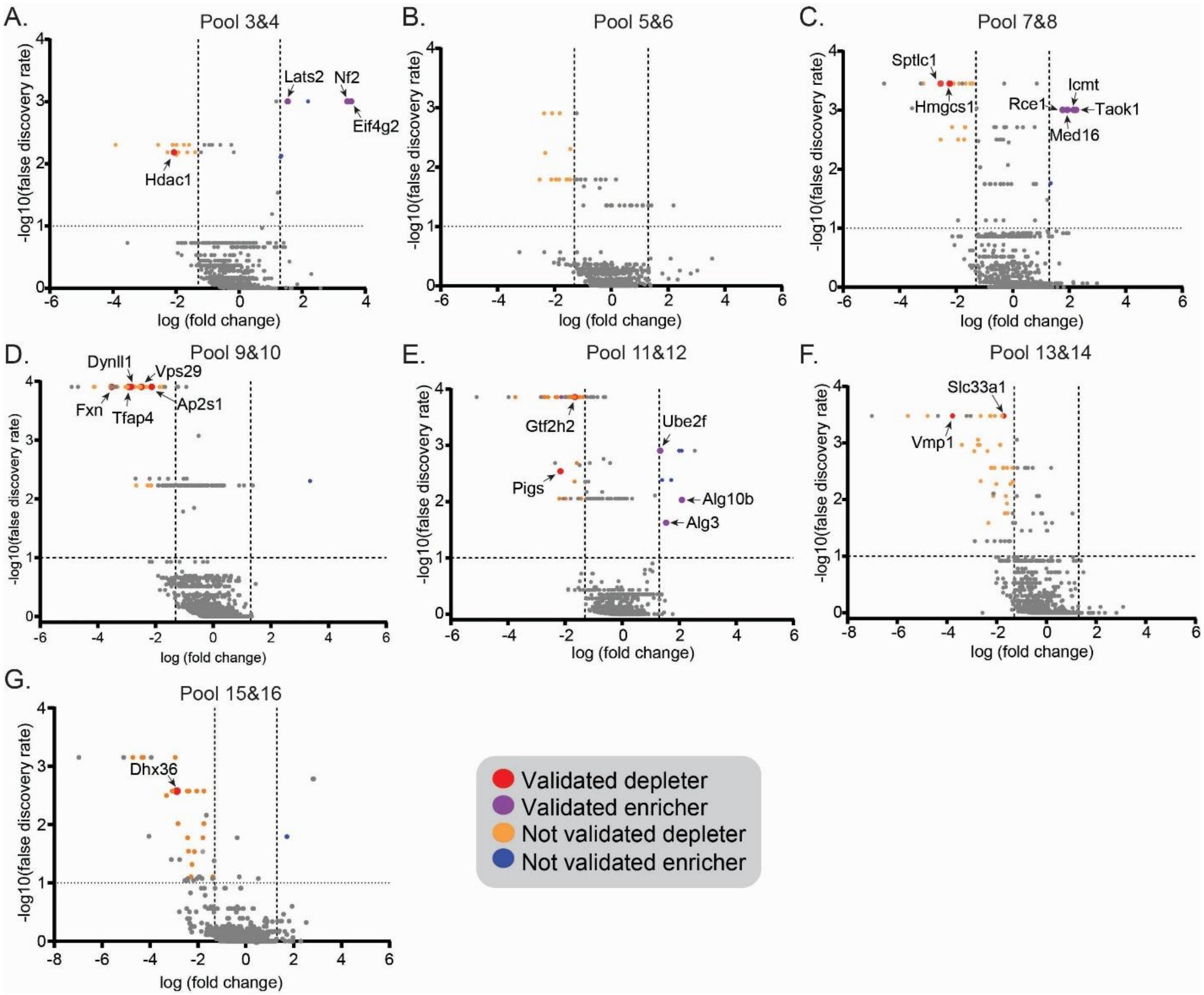


**Figure S6: Iterative in vivo CRISPR-Cas9 screening reveals tumor cell-intrinsic mediators of CAR T cell response.** Volcano plots showing the results of the knock-out screens 2 through 8. “Validated depleter/enricher” refers to genes that were validated in a secondary CRISPR-Cas9 KO screen. “Not validated depleter/enricher” refers to genes that were not confirmed as hits in a secondary CRISPR-Cas9 KO screen. Validated enrichers/depleters are additionally highlighted by name. **A)** Screen 2: Pool 3&4, **B)** Screen 3: Pool 5&6, **C)** Screen 3: Pool 7&8, **D)** Screen 4: Pool 9&10, **E)** Screen 5: Pool 11&12, **F)** Screen 7: Pool 13&14, **G)** Screen 8: Pool 15&16.

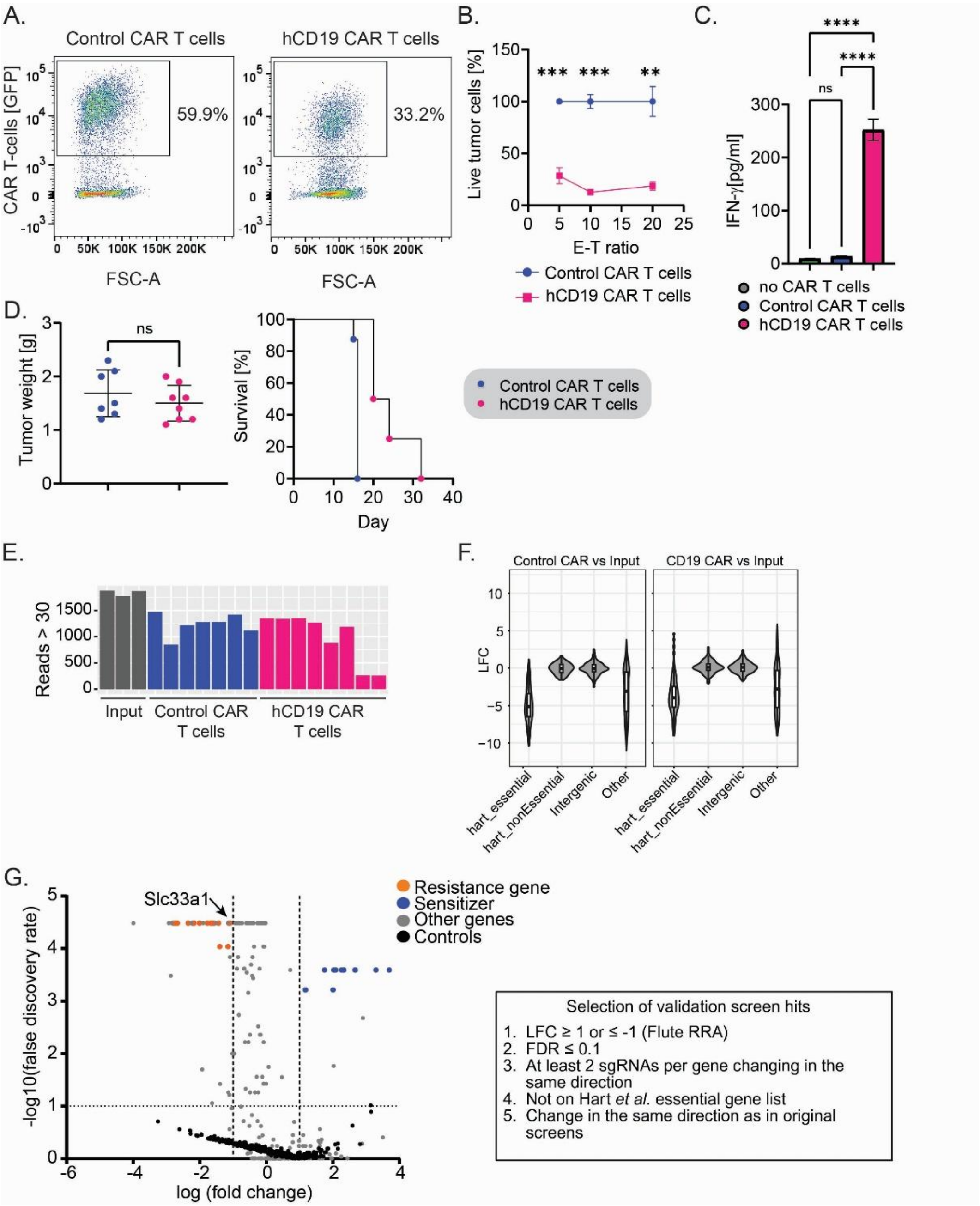


**Figure S7: Validation CRISPR-Cas9 KO screen. A)** A flow cytometry plot showing CAR T cell production efficiency for validation screen. Flow cytometric analysis of CD8+ T cell transduction rates with CAR-GFP constructs. **B)** A graph showing results from a CAR T cell cytotoxicity assay (on the day of *in vivo* injection) involving incubation of hCD19 CAR T cells or control (anti-EGFRvIII) CAR T cells with PDAC cells at multiple effector-to-target ratios (E:T ratios = CAR T cells:PDAC cells = 5:1, 10:1, 20:1) for 24 hours. All conditions were analyzed in triplicate and error bars represent standard deviation. Statistical analysis conducted using unpaired t-test. **C)** A bar graph showing the level of IFN-γ release by hCD19 or control CAR T cells during the CAR T cell assay (E:T ratio = 10:1). All conditions were analyzed in triplicate and error bars represent standard deviation. Statistical analysis conducted using Ordinary One-Way ANOVA (ns > 0.05, * ≤ 0.05, ** p ≤ 0.01, *** p ≤ 0.001, **** p ≤ 0.0001). **D)** Average tumor weights (left) and tumor day harvests (right) as shown in Kaplan-Meier format for the validation CRISPR-Cas9 KO screen. Statistical analysis conducted using Mann-Whitney test. **E)** A bar graph showing the number of represented sgRNA per mouse for each independent CRISPR-Cas9 KO screen. The number of sgRNAs with at least 30 reads is shown for PDAC cells containing the validation CRISPR library on the day of injection (input) compared to tumor cells isolated from mice treated with hCD19 CAR T cells or treated with control CAR T cells. **F)** Violin plots showing the behavior of sgRNAs targeting genes classified as essential or non-essential, as defined by Hart et. al, compared to sgRNAs targeting intergenic regions or all remaining genes in control CAR T cell treated and hCD19 CAR T cell treated mice compared to tumor cells on the day of injection (input). **G)** A volcano plot showing the results of the validation screen. Hits from this screen are split up into "resistance" (orange) and "sensitizer" (blue) genes. To be considered a "hit" each targeted gene had to fulfill five criteria: (1) a LFC of greater than 1 or smaller than -1, (2) a FDR value equal to or smaller than 0.1, (3) at least two out of six guide RNAs changing in the same direction, (4) not considered essential (as defined by Hart et al.), (5) and sgRNAs changed in the same direction as in the original CRISPR-Cas9 KO screens.

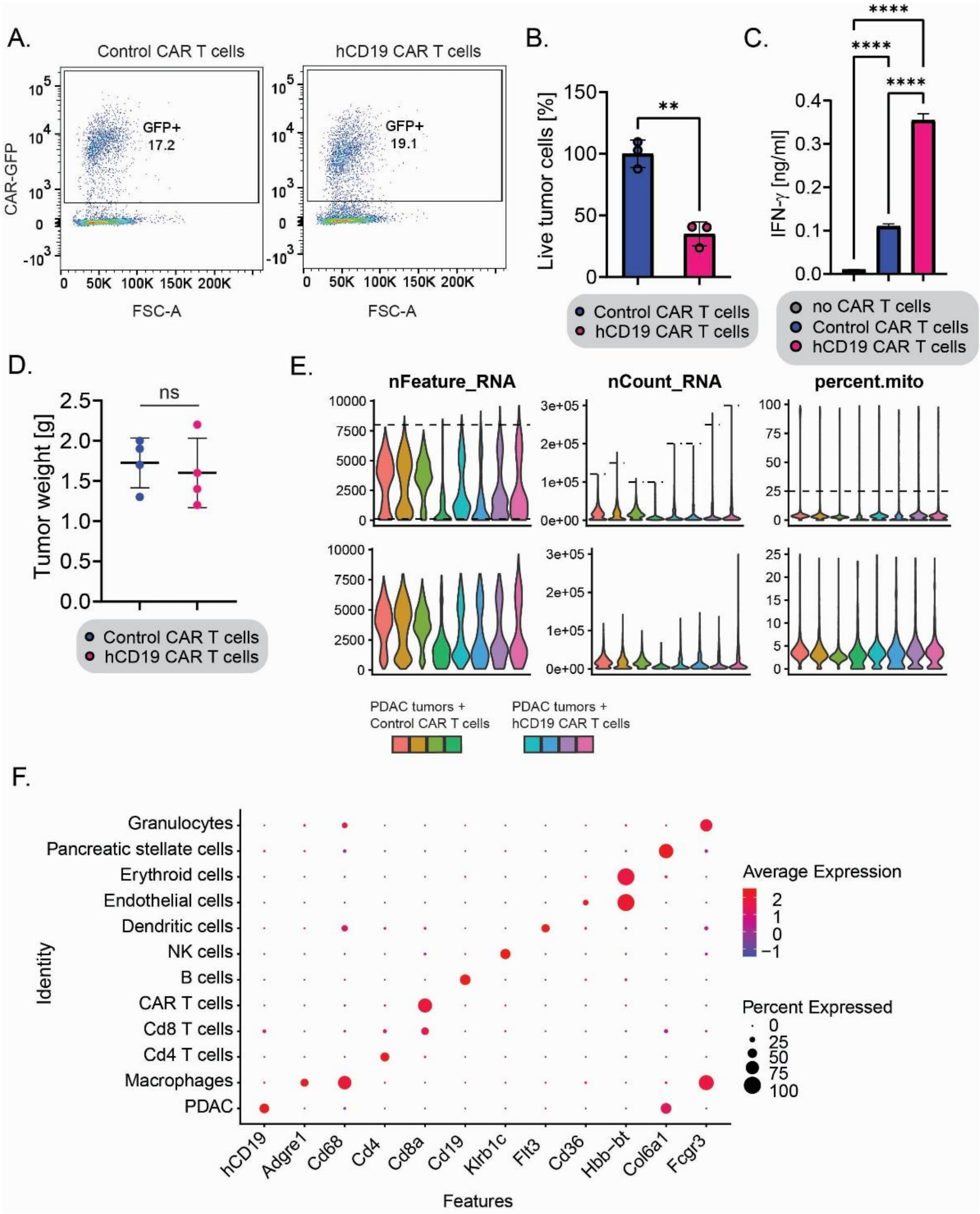


**Figure S8: Single cell RNA sequencing of CAR T cell treated PDAC tumors. A)** Flow cytometric analysis of CD8+ T cell transduction rates with CAR-GFP constructs. **B)** A bar graph showing data from a CAR T cell cytotoxicity assay involving incubation of hCD19 CAR T cells or control (anti-EGFRvIII) CAR T cells with hCD19+ PDAC cells at a 10:1 effector-to-target ratio for 24 hours. All conditions were analyzed in triplicate and error bars represent standard deviation. Statistical analysis conducted using unpaired t-test (ns > 0.05, * ≤ 0.05, ** p ≤ 0.01, *** p ≤ 0.001, **** p ≤ 0.0001). **C)** A bar graph showing the level of IFN-γ release by hCD19 or control CAR T cells during the CAR T cell assay. All conditions were analyzed in triplicate and error bars represent standard deviation. Statistical analysis conducted using Ordinary One-Way ANOVA (ns > 0.05, * ≤ 0.05, ** p ≤ 0.01, *** p ≤ 0.001, **** p ≤ 0.0001). **D)** A graph showing the tumor weight of PDAC tumors treated with control (isolated on day 10) or hCD19 (isolated on day 15) CAR T cells. Tumors are similar in size at the day of isolation. Statistical analysis conducted using t-test (ns > 0.05). **E)** Violin plots showing single cell RNA sequencing data for each PDAC tumor before (top) and after (bottom) filtering of the number of genes detected in each cell **(=**nFeature_RNA >100, <8000), the total number of molecules detected within a cell **(=**nCount_RNA; top 10% removed) and percent mitochondrial reads (=percent.mito; <25%). **F)** Identification of cell clusters in single cell RNA sequencing data. The dot plot is showing the expression levels of multiple genes across tumor cells and stromal cells used for cell type identification.

A.

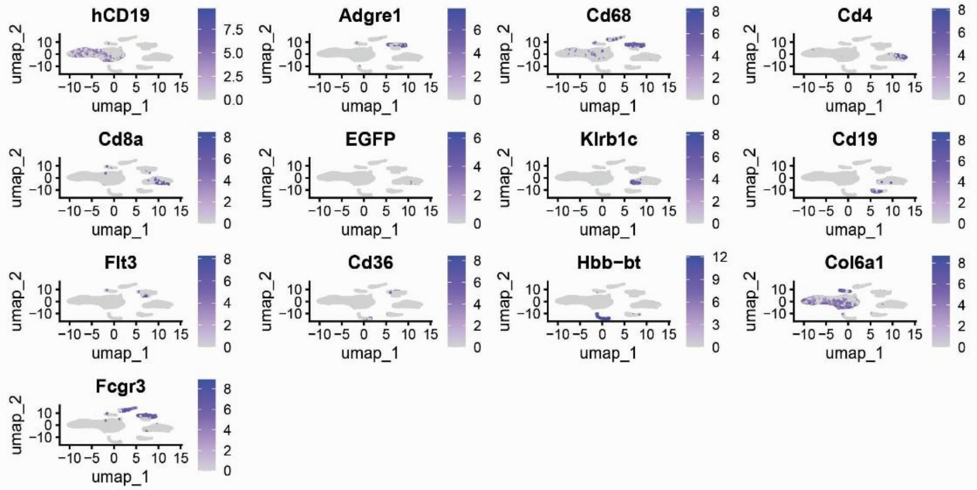


B.

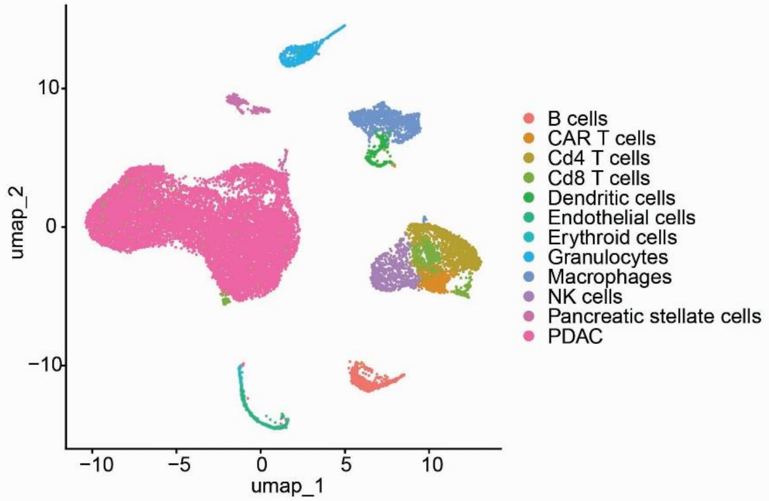


C.

GO terms enriched in hCD19 CAR T cell-treated PDAC tumors

| source | term_name | term_id | adjusted_p_value |
|---|---|---|---|
| GO:MF | extracellular matrix structural constituent | GO:0005201 | 0.0194 |
| REAC | Extracellular matrix organization | REAC:R-HSA-1474244 | 0.0001 |
| REAC | Non-integrin membrane-ECM interactions | REAC:R-HSA-3000171 | 0.0035 |
| WP | Wnt beta catenin signaling in leukemia | WP:WP3658 | 0.0385 |
| WP | Acute myeloid leukemia | WP:WP5293 | 0.0477 |

**Figure S9: Single cell RNA sequencing cluster annotation and gene ontology. A)** Feature plots showing the expression of different genes across all clusters of cells isolated from PDAC tumors in a UMAP. These are examples of genes that were used for the identification and annotation of the shown cell clusters. **B)** A UMAP showing the annotated tumor and stromal cell clusters isolated from PDAC tumors. **C)** Gene ontology (GO) analysis of top 100 enriched genes in hCD19 CAR T cell treated tumors compared to control CAR T cell treated tumors.

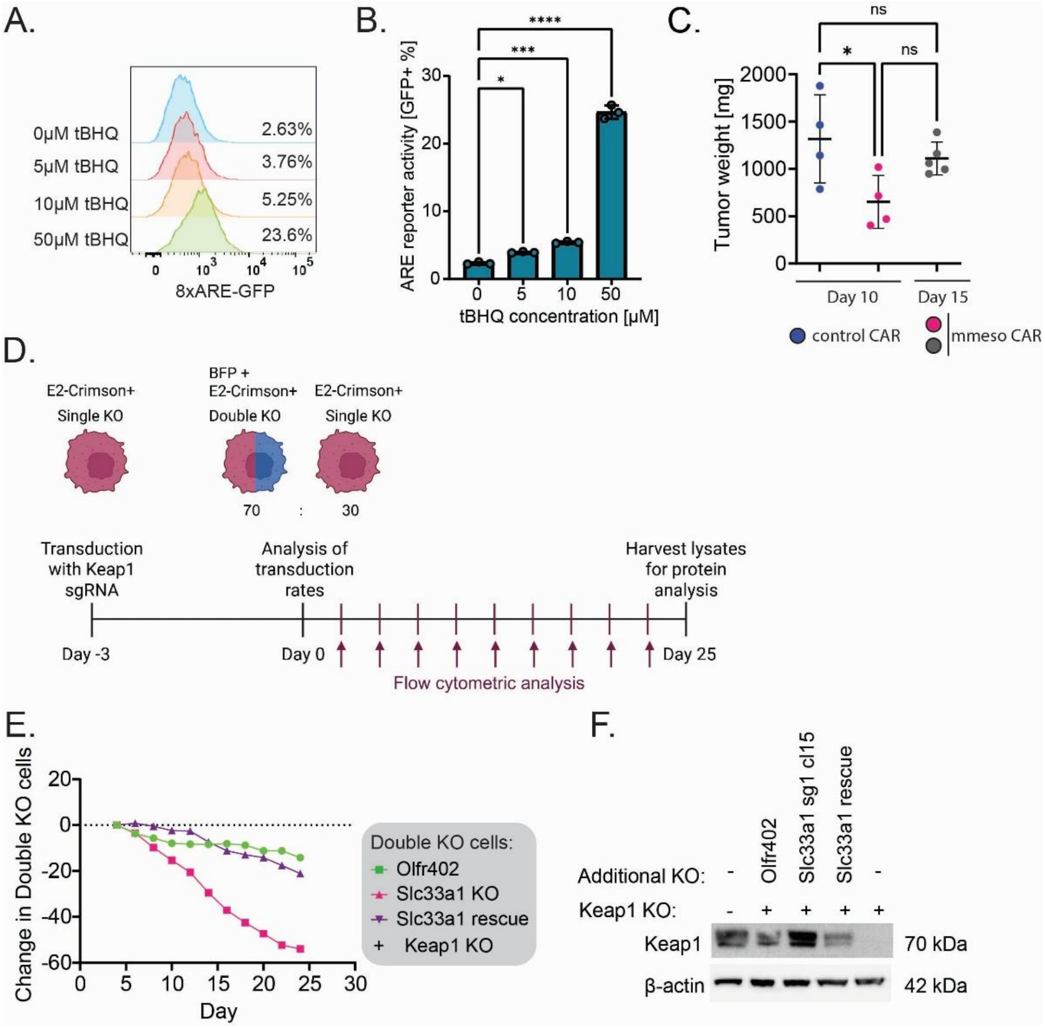


**Figure S10: *Nrf2*-activity in PDAC cells. A)** Flow cytometry showing *Nrf2* reporter expression. Treatment of WT PDAC cells expressing an *NRF2* 8xARE-GFP reporter with the Nrf2 activator tBHQ for 24 hours *in vitro* leads to dose-dependent expression of GFP as measured by flow cytometry. **B)** A bar graph showing quantification of *NRF2* 8xARE-GFP reporter activity (as % GFP) after treatment with different doses of tBHQ for 24 hours. All conditions were analyzed in triplicate and error bars represent standard deviation. Statistical analysis conducted using Ordinary One-Way ANOVA (ns > 0.05, * ≤ 0.05, ** p ≤ 0.01, *** p ≤ 0.001, **** p ≤ 0.0001). **C)** A graph showing the weight of PDAC tumors formed by WT PDAC cells expressing an *NRF2* 8xARE-GFP reporter in C57BL/6 mice 10 days and 15 days after tumor cell injection. Statistical analysis with Ordinary One-Way ANOVA test (ns > 0.05, * ≤ 0.05, ** p ≤ 0.01, *** p ≤ 0.001, **** p ≤ 0.0001). **D)** Analysis of the effect of Keap1 KO on the proliferation of control (*Olfr402*) KO, *Slc33a1* KO or *Slc33a1* re-expression (rescue) *in vitro*. E2-Crimson+ PDAC cells with control KO, Slc33a1 KO or Slc33a1 rescue were transduced with a previously validated Keap1 sgRNA (BFP+). Three days after transduction, the transduction rates were analyzed by flow cytometry. Single KO and double KO cells were at an approximately 30:70 ratio at the beginning of the long-term co-culture. Throughout the co-culture, cells were analyzed every other day by flow cytometry. On day 25, lysates were harvested for protein analysis. **E)** A graph showing growth dynamics of Keap1 KO PDAC cells in a control (*Olfr402*) KO, *Slc33a1* KO or *Slc33a1* re-expression (rescue) background *in vitro*. The percentages of double KO cells were compared to the percentages of single KO cells over 25 days. *Keap1* and *Slc33a1* double KO cells get depleted over time. **F)** A western blot showing Keap1 protein expression at the endpoint of long-term *in vitro* co-culture from (E). In the *Slc33a1* KO condition, *Slc33a1/Keap1* double-KO cells are depleted.

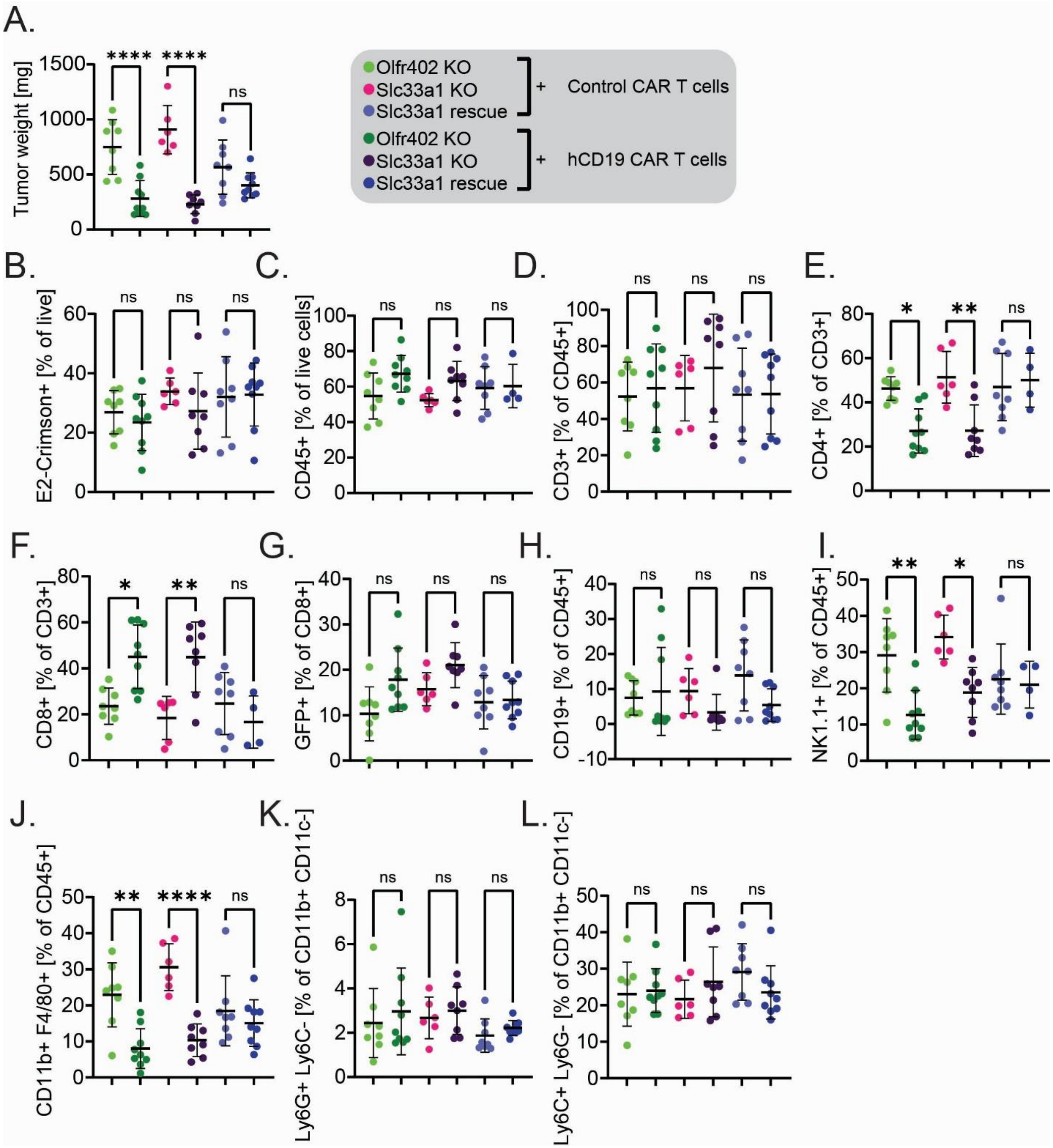


**Figure S11: Analysis of *Slc33a1* KO tumor composition by flow cytometry. A)** A graph showing the tumor weight of PDAC tumors with a control (*Olfr402*) KO, *Slc33a1* KO, or re-expression of *Slc33a1* (rescue), that were treated with control or hCD19 CAR T cells. All tumors were isolated on day 10 after tumor cell injection. PDAC tumors with *Slc33a1* or control KO treated with hCD19 CAR T cells are significantly smaller than control (EGFRvIII) CAR T cell treated tumors, but not *Slc33a1* rescue tumors. **B-L) Analysis of PDAC tumor composition by flow cytometry. B)** Percentage of E2-Crimson+ PDAC cells of all live cells. **C)** Percentage of CD45+ immune cells of all live cells. **D)** Percentage of CD3+ T cells of all immune cells. **E)** Percentage of CD4+ T cells of CD3+ T cells. CD4+ T cells are significantly reduced in PDAC tumors with control KO or *Slc33a1* KO treated with hCD19 CAR T cells compared to control CAR T cell treatment. **F)** Percentage of CD8+ T cells of CD3+ T cells. CD8+ T cells are significantly enriched in

PDAC tumors with control KO or *Slc33a1* KO treated with hCD19 CAR T cells compared to control CAR T cell treatment. **G)** GFP+ CAR T cells as a percentage of all CD8 T cells. **H)** Percentage of CD19+ B cells of all immune cells. **I)** Percentage of NK1.1+ NK cells of all immune cells. **J)** Percentage of CD11b+ F4/80+ Macrophages of all immune cells. **K)** Percentage of Ly6G+ Ly6C- Neutrophils of all CD11b+ CD11c- immune cells. **L)** Percentage of Ly6C+ Ly6G- MDSCs of all CD11b+ CD11c- immune cells.

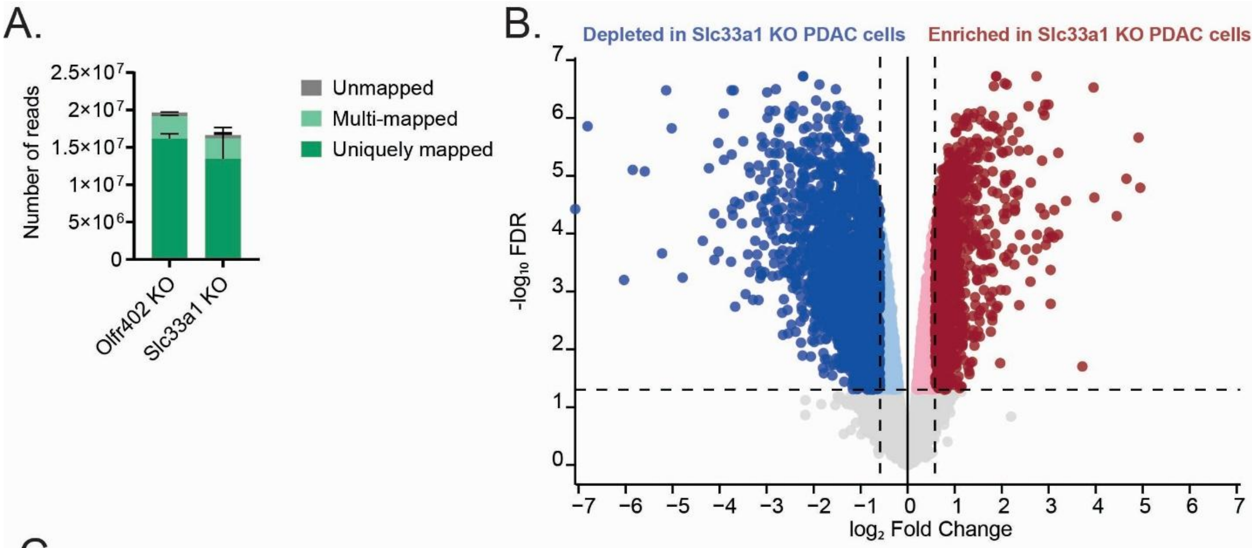


C.

Depleted in Slc33a1 KO PDAC cells

| MSigDB Hallmark term | Size | NES | NOM p-value | FDR q-value |
|---|---|---|---|---|
| Interferon Gamma Response | 102 | -2.8781 | 0 | 0 |
| Interferon Alpha Response | 62 | -2.835 | 0 | 0 |
| IL6 JAK STAT3 Signaling | 43 | -2.2432 | 0 | 0 |
| Allograft Rejection | 57 | -2.1142 | 0 | 0 |
| Complement | 67 | -2.0055 | 0 | 0 |
| Epithelial Mesenchymal Transition | 60 | -1.9226 | 0 | 0 |
| Angiogenesis | 10 | -1.9174 | 0 | 0.000151143 |
| Inflammatory Response | 58 | -1.893 | 0 | 0.000396751 |
| Coagulation | 36 | -1.8159 | 0 | 0.000940448 |
| Apical Junction | 56 | -1.777 | 0 | 0.001481205 |

Enriched in Slc33a1 KO PDAC cells

| MSigDB Hallmark term | Size | NES | NOM p-value | FDR q-value |
|---|---|---|---|---|
| MYC Targets V1 | 145 | 2.96 | 0 | 0 |
| Unfolded Protein Response | 64 | 2.71872 | 0 | 0 |
| MYC Targets V2 | 33 | 2.46046 | 0 | 0 |
| E2F Targets | 113 | 2.39803 | 0 | 0 |
| MTORC1 Signaling | 79 | 2.19919 | 0 | 0 |
| Oxidative Phosphorylation | 82 | 2.10713 | 0 | 0.000149276 |
| DNA Repair | 64 | 1.96955 | 0 | 0.000127951 |
| G2M Checkpoint | 80 | 1.8049 | 0 | 0.001343484 |
| Adipogenesis | 57 | 1.44245 | 0.003154574 | 0.027964373 |
| Fatty Acid Metabolism | 31 | 1.39581 | 0.014409222 | 0.038154948 |

**Figure S12: Bulk tumor cell transcriptome profiling of *Slc33a1* KO PDAC cells. A)** Bulk RNA sequencing read assignment for Control (*Olfr402*) or *Slc33a1* KO PDAC cells isolated from orthotopic tumors. The majority of reads are uniquely mapped. **B)** Volcano plot showing the differentially expressed genes of *Slc33a1* KO PDAC cells compared to control PDAC cells. Genes depicted in blue are depleted in *Slc33a1* KO cells compared to control KO cells. Genes depicted in red are enriched in *Slc33a1* KO cells compared to control KO cells. The cutoff for gene significance was set at a log fold change of |1.5| and an adjusted p-value of < 0.05. The y-axis depicts the -$\log_{10}$ FDR and the x-axis depicts the $\log_2$ fold change. **C)** GSEA analysis on differentially expressed genes in *Slc33a1* KO cells compared to control KO cells using the MSigDB Hallmark gene set. The top 10 pathways enriched (red) or depleted (blue) in *Slc33a1* KO cells compared to control KO cells are shown.

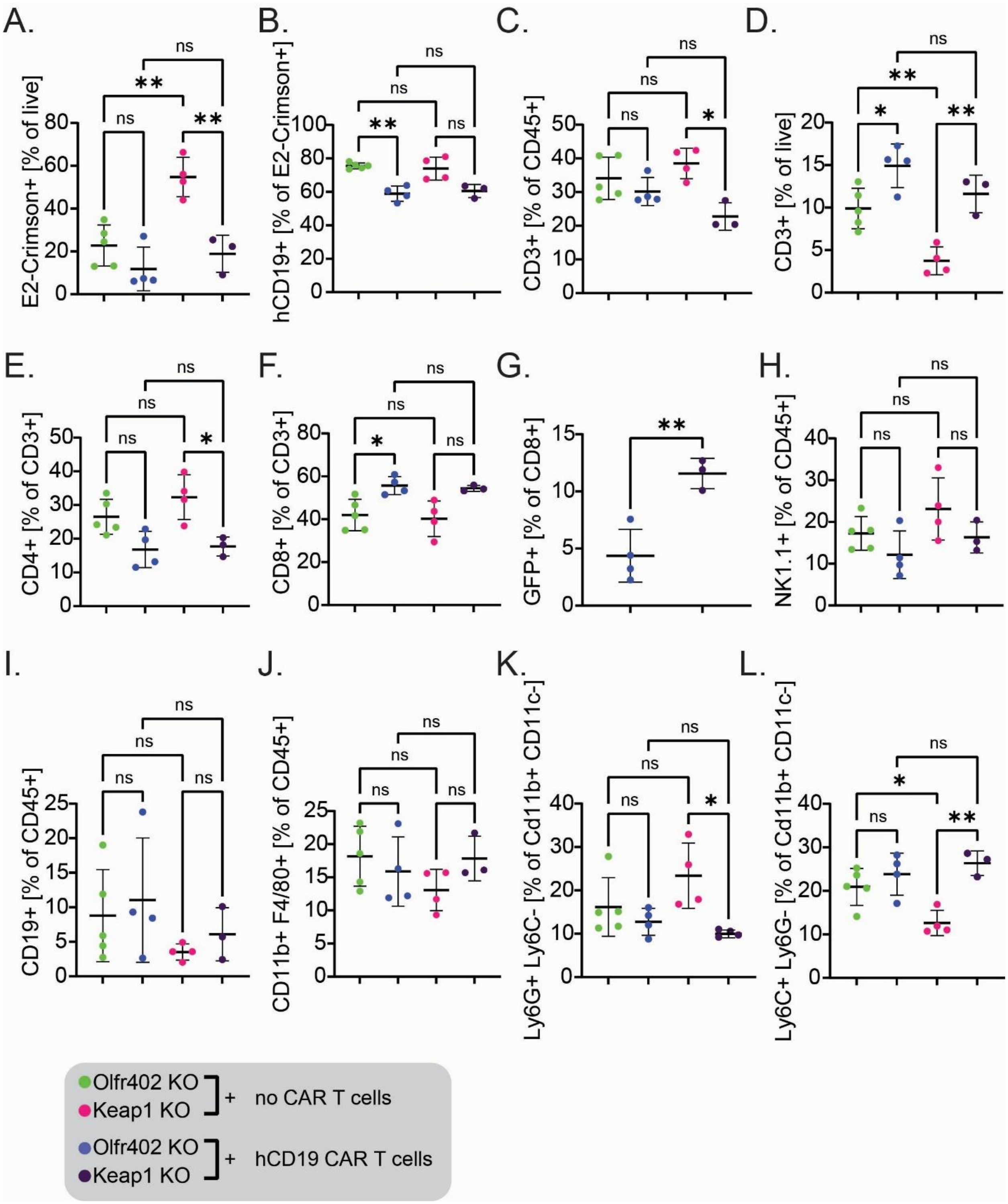


**Figure S13: Analysis of *Keap1* KO tumor composition by flow cytometry. A)** Percentage of E2-Crimson+ PDAC cells of all live cells. Untreated Keap1 KO tumors contain a significant higher proportion of PDAC cells. **B)** Percentage of antigen-positive (hCD19+) PDAC cells. **C)** Percentage of CD3+ T cells of all immune cells. **D)** Percentage of CD3+ T cells of all live cells. CD3+ tumor cells are enriched in hCD19 CAR T cell treated PDAC tumors . **E)** Percentage of CD4+ T cells of CD3+ T cells. **F)** Percentage of CD8+ T cells of CD3+ T cells. **G)** GFP+ CAR T cells as a percentage of all CD8 T cells. **H)** Percentage of NK1.1+

NK cells of all immune cells. **I)** Percentage of CD19+ B cells of all immune cells. **J)** Percentage of CD11b+ F4/80+ Macrophages of all immune cells. **K)** Percentage of Ly6G+ Ly6C- Neutrophils of all CD11b+ CD11c- immune cells. **L)** Percentage of Ly6C+ Ly6G- MDSCs of all CD11b+ CD11c- immune cells.

A.

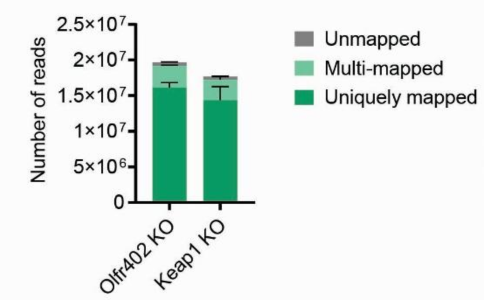


B.

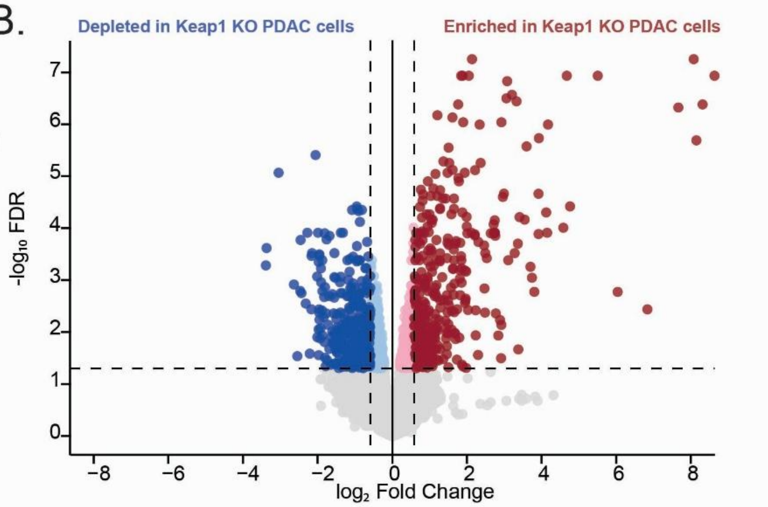


C.

**Depleted in Keap1 KO PDAC cells**

| MSigDB Hallmark term | Size | NES | NOM p-value | FDR q-value |
|---|---|---|---|---|
| Interferon Gamma Response | 109 | -2.445 | 0 | 0 |
| Interferon Alpha Response | 54 | -2.436 | 0 | 0 |
| Inflammatory Response | 83 | -2.109 | 0 | 0 |
| TNFA Signaling VIA NFKB | 82 | -2.106 | 0 | 0 |
| Epithelial Mesenchymal Transition | 63 | -2.026 | 0 | 0 |
| Coagulation | 35 | -1.966 | 0 | 0 |
| KRAS Signaling UP | 73 | -1.874 | 0 | 0.000310041 |
| Allograft Rejection | 65 | -1.853 | 0 | 0.000542571 |
| IL6 JAK STAT3 Signaling | 38 | -1.833 | 0 | 0.000482285 |
| Hypoxia | 61 | -1.829 | 0 | 0.000434057 |

**Enriched in Keap1 KO PDAC cells**

| MSigDB Hallmark term | Size | NES | NOM p-value | FDR q-value |
|---|---|---|---|---|
| Xenobiotic Metabolism | 36 | 2.52093 | 0 | 0 |
| Reactive Oxygen Species Pathway | 20 | 2.35896 | 0 | 0 |
| Oxidative Phosphorylation | 91 | 2.14969 | 0 | 0 |
| MYC Targets V1 | 81 | 2.08381 | 0 | 0 |
| MYC Targets V2 | 38 | 2.02866 | 0 | 0 |
| Adipogenesis | 61 | 1.94714 | 0 | 0 |
| Fatty Acid Metabolism | 47 | 1.69095 | 0 | 0.00473996 |
| Bile Acid Metabolism | 10 | 1.62308 | 0.005791506 | 0.00979263 |
| Peroxisome | 13 | 1.58617 | 0.005813953 | 0.0124936 |
| MTORC1 Signaling | 29 | 1.49458 | 0.001828154 | 0.02866359 |

D.

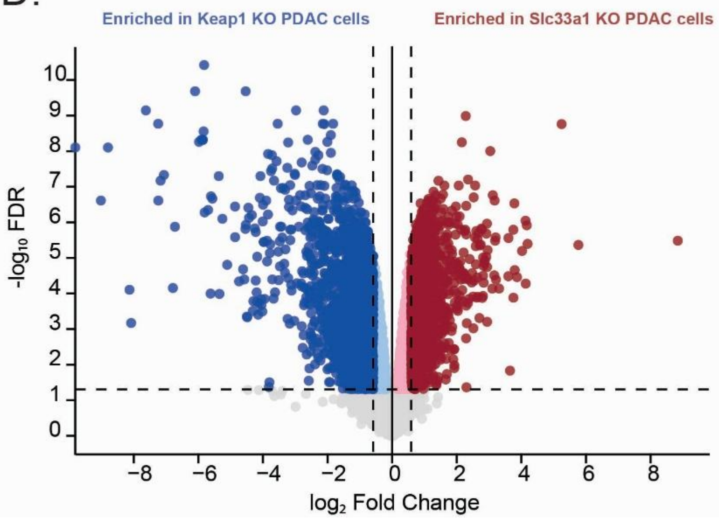


E.

| MSigDB Hallmark term | Size | NES | NOM p-value | FDR q-value |
|---|---|---|---|---|
| MYC Targets V1 | 99 | 2.66034 | 0 | 0 |
| Unfolded Protein Response | 67 | 2.46557 | 0 | 0 |
| E2F Targets | 90 | 2.32497 | 0 | 0 |
| MYC Targets V2 | 38 | 2.16408 | 0 | 0 |
| G2M Checkpoint | 61 | 2.06507 | 0 | 0 |
| MTORC1 Signaling | 67 | 1.91177 | 0 | 0.000453972 |
| DNA Repair | 53 | 1.52754 | 0 | 0.018288597 |
| TNFA Signaling VIA NFKB | 46 | 1.38732 | 0.002695418 | 0.056860025 |
| PI3K AKT MTOR Signaling | 29 | 1.31055 | 0.049261084 | 0.098259773 |
| Oxidative Phosphorylation | 52 | 1.30716 | 0.018018018 | 0.09097604 |

| MSigDB Hallmark term | Size | NES | NOM p-value | FDR q-value |
|---|---|---|---|---|
| Reactive Oxygen Species Pathway | 18 | -2.1249 | 0 | 0 |
| Xenobiotic Metabolism | 54 | -1.9925 | 0 | 0 |
| Angiogenesis | 12 | -1.9643 | 0 | 0 |
| Apical Surface | 13 | -1.8718 | 0 | 0.00079536 |
| Bile Acid Metabolism | 28 | -1.8496 | 0 | 0.001060481 |
| Interferon Gamma Response | 79 | -1.6151 | 0 | 0.017851422 |
| Peroxisome | 28 | -1.5794 | 0.004975124 | 0.021058113 |
| Interferon Alpha Response | 49 | -1.506 | 0.001680672 | 0.033935377 |
| KRAS Signaling DN | 27 | -1.4403 | 0.026315789 | 0.057619442 |
| Epithelial Mesenchymal Transition | 52 | -1.4266 | 0.007974482 | 0.060023198 |

**Figure S14: Bulk tumor cell transcriptome profiling of *Keap1* KO PDAC cells. A)** Bulk RNA sequencing read assignment for Control (*Olfr402*) or *Keap1* KO PDAC cells isolated from orthotopic tumors. The majority of reads are uniquely mapped. **B)** Volcano plot showing the differentially expressed genes of *Keap1* KO PDAC cells compared to control PDAC cells. Genes depicted in blue are depleted in *Keap1* KO cells compared to control KO cells. Genes depicted in red are enriched in *Keap1* KO cells compared to control KO cells. The cutoff for gene significance was set at a log fold change of |1.5| and an adjusted p-value of < 0.05. The y-axis depicts the $-\log_{10}$ FDR and the x-axis depicts the $\log_2$ fold change. **C)** GSEA analysis on differentially expressed genes in *Keap1* KO cells compared to control KO cells using the MSigDB Hallmark gene set. The top 10 pathways enriched (red) or depleted (blue) in *Keap1* KO cells compared to control KO cells are shown. **D)** Volcano plot showing the differentially expressed genes of *Keap1* KO PDAC cells compared to *Slc33a1* KO PDAC cells. Genes depicted in blue are enriched in *Keap1* KO cells compared to *Slc33a1* KO cells. Genes depicted in red are enriched in *Slc33a1* KO tumor cells compared to *Keap1* KO cells. The cutoff for gene significance was set at a log fold change of |1.5| and an adjusted p-value of < 0.05. The y-axis depicts the $-\log_{10}$ FDR and the x-axis depicts the $\log_2$ fold change. **E)** GSEA analysis on differentially expressed genes in *Keap1* KO cells compared to *Slc33a1* KO cells using the MSigDB Hallmark gene set. The top 10 pathways enriched in Slc33a1 KO cells (red) or enriched in *Keap1* KO cells (blue) are shown.

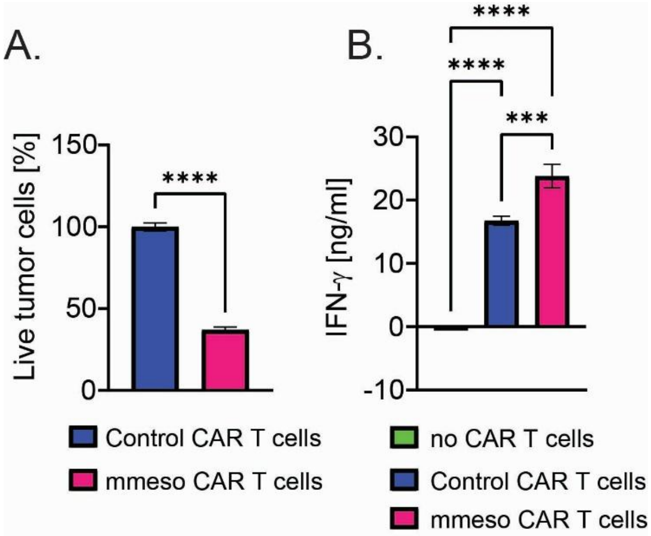


**Figure S15: WT PDAC cells are efficiently killed by mmeso CAR T cells. A)** A bar graph showing CAR T cell killing after incubation of mmeso CAR T cells or control (EGFRvIII) CAR T cells with PDAC cells (E:T ratio = 10:1) for 24 hours. All conditions were analyzed in triplicate. Error bars represent standard deviation. Statistical analysis with Ordinary One-Way ANOVA test (ns > 0.05, * ≤ 0.05, ** p ≤ 0.01, *** p ≤ 0.001, **** p ≤ 0.0001). **B)** A bar graph showing the average IFN-γ release by mmeso or control (EGFRvIII) CAR T cells during the CAR T cell cytotoxicity assays. All conditions were analyzed in triplicate and error bars represent standard deviation. Statistical analysis conducted using Ordinary One-Way ANOVA test (ns > 0.05, * ≤ 0.05, ** p ≤ 0.01, *** p ≤ 0.001, **** p ≤ 0.0001).
